# Estimating population split times and migration rates from historical effective population sizes

**DOI:** 10.1101/2022.06.17.496540

**Authors:** Vladimir Shchur, Débora Y. C. Brandt, Anna Ilina, Rasmus Nielsen

## Abstract

The estimation of effective population sizes (*N_e_*) through time is of fundamental interest in population genetics, but the interpretation of *N_e_* as the effective number of breeding individuals in the population is challenged by the effect of population structure. In fact, variation in *N_e_* reported in many studies may be a consequence of changes in migration rates between populations rather than changes in actual population size. We address this long-standing problem here by constructing joint models of population size changes, migration, and divergence that can adjust temporal estimates of *N_e_* and estimate the actual *N_e_* of a local deme connected to another population through migration. We also develop a method for estimating divergence times and migration rates taking into account complex scenarios of changing population sizes. We apply the method to previously published data from humans, and show that, when taking migration and changes in *N_e_* into account, the estimated divergence between the San and Dinka populations is approximately 108 kya, and not 255 kya as reported in a previous study. Using simulations, we demonstrate that the previously reported and surprisingly old estimates of divergence between San and Dinka is in fact caused by a quantifiable estimation bias due to changes in *N_e_* through time.

## 1 Introduction

Effective population size is one of the main characteristics of the demography of a population, and an important parameter determining evolutionary processes. Although it is related to census population sizes, it does not correspond to it in most cases. For example, for humans, effective population size is estimated to be on the order of ten thousand, which is much smaller than actual population size, in the order of billions [Henn et al., 2012].

Effective population size has been formally defined in many different ways. The concept was originally introduced by Wright [1931] in the context of describing causes of random variation in allele frequencies through time (i.e. genetic drift). In that context, effective population size is “*the number of individuals in a theoretically ideal population having the same magnitude of **random genetic drift** as the actual population*” [Hartl and Clark, 2007]. Common models for this ideal population are standard Wright-Fisher [Fisher, 1930, Wright, 1931] or Moran [Moran, 1958] models. In an actual population, a combination of population size, inbreeding, unequal sex ratios and variance in offspring number all contribute to genetic drift, and thus to the effective population size. In population models where these parameter values are known, effective population size can be calculated directly using well-established equations [Hartl and Clark, 2007].

In practice, because effective population size measures genetic drift, and drift determines the amount of genetic diversity in the idealized population, effective population size is often interpreted as a measure of genetic diversity. The definition of effective population size then changes slightly to *the number of individuals in a theoretically ideal population having the same amount of **genetic variation** as the actual population*. Under this definition, effective population size can be calculated directly from measures of genetic diversity of real populations. In a real population, however, the level of genetic diversity is affected by processes other than drift, such as migration and natural selection. When the effective population size is estimated from genetic diversity, it is affected by these other processes. In other words, when we define effective population size as a measure of genetic diversity, it no longer reflects only drift.

The Wright-Fisher and Moran population models mentioned earlier, as well as many others, converge to Kingman’s coalescent model when population sizes are large [Kingman, 1982, Sjödin et al., 2005, Wakeley and Sargsyan, 2009]. In this model, effective population size is a parameter determining the probability distribution for coalescence times (time to a common ancestor) of two lineages. More specifically, the **coalescent** effective population size is *the inverse of the coalescence rate*. Because coalescence times predict most measures of genetic diversity in a population, the coalescent effective population size has been argued to be the most general definition of effective population size [Sjödin et al., 2005], and it is the definition we will use here.

### 1.1 Methods to estimate historical effective population size

Many methods were developed to estimate the variation of effective population size through time. The most widely used one is PSMC [Li and Durbin, 2011], which uses a single diploid genome (or a pair of haploid genomes) to infer past effective population sizes. PSMC is based on a sequentially Markovian coalescent model [McVean and Cardin, 2005], where states (coalescence times, discretized) change along a DNA sequence. A Hidden Markov Model (HMM) approach is used to infer the distribution of coalescent times between a pair of lineages. The emission probabilities of this HMM are the probabilities of observing a site that is variable (heterozygous) or non-variable (homozygous), given a coalescence time. The underlying intuition is that the probability of observing a heterozygous site increases with coalescence time. The effective population size is then given by the inverse of the coalescence rate. There are many other HMM-based methods for inference of historical effective population size: coalHMM [Hobolth et al., 2007, Dutheil et al., 2009], diCal [Sheehan et al., 2013], MSMC [Schiffels and Durbin, 2014], SMC++ [Terhorst et al., 2017] (see Spence et al. [2018] for a review).

Previous studies have shown how these and other methods can infer past effective population sizes that are very different from simulated census population sizes when populations are structured [Heller et al., 2013, Mazet et al., 2016, Chikhi et al., 2018]. These studies showed how population structure can lead to misinterpretation of past effective population size plots. While it is true that PSMC plots can be often misinterpreted, we argue that the change in past effective population size due to population structure is an expected and desirable behaviour, since these methods infer the coalescent effective population size, and not the census population sizes.

Migration, for example, is a phenomenon that generally increases the coalescent effective population size of the population receiving migrants, since incoming migrants will likely increase genetic diversity. Interestingly, if we consider the alternative definition of effective population size as the change in allele frequency through time due to drift, the effect of migration on effective population size is the opposite: migration introduces sudden changes in allele frequency, which can be interpreted as strong drift, and thus small effective population size [Wang and Whitlock, 2003]. This effect is expected only in the short term, and it is reversed in the longer term as populations approach an equilibrium [Wang and Whitlock, 2003]. Here, however, we are focusing on the coalescent effective population size, which tends to increase with immigration. We emphasize that the effective population size inferred with PSMC should be interpreted as the amount of genetic variation in a population through time. Therefore, PSMC results are informative about both population size and migration. Nonetheless, as we will show, inferences of effective population size from PSMC can in some cases be biased when the transition probabilities of the HMM underlying PSMC inferences cannot adequately fit the true transitions in coalescence times along the genome in the presence of migration.

In this work we make an effort to disentangle the effects of migration from effective population sizes. We discuss the case of two fully exchangeable populations (*e.g*. Wright-Fisher populations) with migration between them. A sample from any of the two admixed populations contains footprints of historical effective population size of both parental populations. We formalize the concept of *local* effective population size, which is the effective population size of the parental populations, after accounting for the effect of migration. We show that the local effective population size can be determined from the the ordinary effective population size estimated by PSMC if migration rates are known or inferred.

We also develop a method called MiSTI (for “migration and split time inference”), which infers split time and migration rates under a model of two populations that exchange migrants after their split from a common ancestor. To do so, MiSTI combines information from the joint site frequency spectrum of two diploid samples with the ordinary historical effective population sizes (as inferred by PSMC). MiSTI also uses the inferred migration rates to recover the local effective population size, *i.e*. to “correct” the PSMC curves for the effect of admixture. By applying this method to simulated data we show scenarios where PSMC finds a good approximation of the simulated effective population size, and scenarios where PSMC results do not correspond to the true effective population size. We also show that MiSTI appropriately corrects PSMC curves for the effect of migration, when migration rates are known and PSMC estimates are close to the true effective population size. Next, we apply MiSTI to data from humans and show i) How MiSTI can correct the effect of Neanderthal admixture on the historical effective population size of a human genome of European ancestry (CEU) and ii) What split times and migration rates best fit a model of split time and migration between pairs of human populations.

### 1.2 Other methods that infer population split times and/or migration rates from a pair of diploid genomes

Wang et al. [2020] developed a method, MSMC-IM, that also infers migration rates from historical effective population sizes. MSMC-IM fits an isolation-migration model with continuous symmetric migration to the inverse coalescence rates inferred by MSMC2. Instead of explicitly modelling a split time point, they model population split as a continuous process. The event of two populations merging backwards in time is represented as an increase in migration rates, to a point where both populations exchange migrants freely. In contrast to MiSTI, MSMC-IM does not aim to recover the local effective population size. Other differences worth pointing out are that MiSTI allows for asymmetric migration between populations and it uses PSMC instead of MSMC, which requires phased genomes.

Song et al. [2017] also tackled the problem of fitting an isolation-migration model to infer population split times from PSMC results. Their approach differs from ours methodologically and conceptually. In terms of methodology, Song et al. [2017] use an ABC approach to fit parameters, while we compute the composite likelihood of parameter values based on equations derived analytically. Conceptually, we formalize the distinction between the ordinary effective population size of admixed samples (often inflated by migration) and the local effective population size of its parental populations, which we can recover in the presence of migration, while Song et al. [2017] does not make this distinction.

Arredondo et al. [2021] use yet another approach. Their method, SNIF (Structured Non-stationary Inferential Framework) fits the curves of coalescent effective population size through time (which they denominate inverse instantaneous coalescence rate, IICR), as inferred by PSMC, to island models with symmetric migration and constant deme size. This method allows to infer the number of demes and migration rates among demes from IICR curves alone.

Schlebusch et al. [2017] introduced the TT-method [Sjödin et al., 2021], which can also infer population split times from two diploid genomes representing each of the two populations. The TT-method uses the joint site frequency spectrum of these two genomes to analytically calculate split times. It relies on two assumptions that are relaxed in MiSTI: 1) the effective population size of the ancestral population remains constant and 2) there is no migration between populations after the split. We compared MiSTI and TT-method inferences of split times between human populations, and we show through simulations that the first assumption of TT-method leads to large errors in the inferred split times for historical effective population sizes similar to those of human populations, even in the absence of migration.

## 2 Methods

### 2.1 Historical effective population size

As previously discussed, effective population size can be defined as the average time to coalescence of two lineages, measured in number of generations [Wakeley and Sargsyan, 2009]. Under the standard coalescence model with a single homogeneous population [Kingman, 1982], the interpretation is simple. For an effective population size *N* >> 1 the rate of coalescence is *λ* = 1/*N* per generation, and the expected waiting time to coalescence is *λ*^-1^ = *N* generations. This definition can be naturally extended in order to define the historical effective population size. Consider the coalescence rate at time *t, λ*(*t*), between the pairs of lineages from a population. The time *t* = 0 corresponds to the present and *t* increases toward the past. The coalescent rate *λ*(*t*) determines an inhomogenous Poisson process which describes the distribution of coalescent times. Hence, the probability distribution of coalescent times *T_c_* is

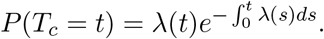

We define the inverse of *λ*(*t*) as the ordinary *historical effective population size*

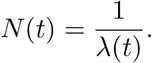

This quantity depends on population structure and demography, and it is a parameter which allows mapping of a real population with complex demography and structure on a single idealized population (e.g., a Wright-Fisher population) that is similar with respect to some property, as described in the Introduction. We note that this concept of historical effective population size is useful for interpreting the results of methods such as PSMC that allow inferences of varying effective population size through time [Li and Durbin, 2011, Spence et al., 2018]. We will show that though for some scenarios PSMC indeed infers a good estimate of the ordinary effective population size, in other scenarios with migration, PSMC infers a biased estimate of effective population size.

Using standard population genetic theory, it is possible to explore the effect of population structure on effective population size [Mazet et al., 2016, Chikhi et al., 2018]. Assume, for example, that an observed (modern) population *S_m_* is formed by admixture of several parental populations. To determine its historical effective population size, we need to trace pairs of lineages from *S_m_* back in time, until their coalescence. The lineages can switch between parental populations (Fig. 1), hence the genetic variation of *S_m_* has a footprint from each of these populations.

**Figure 1:**
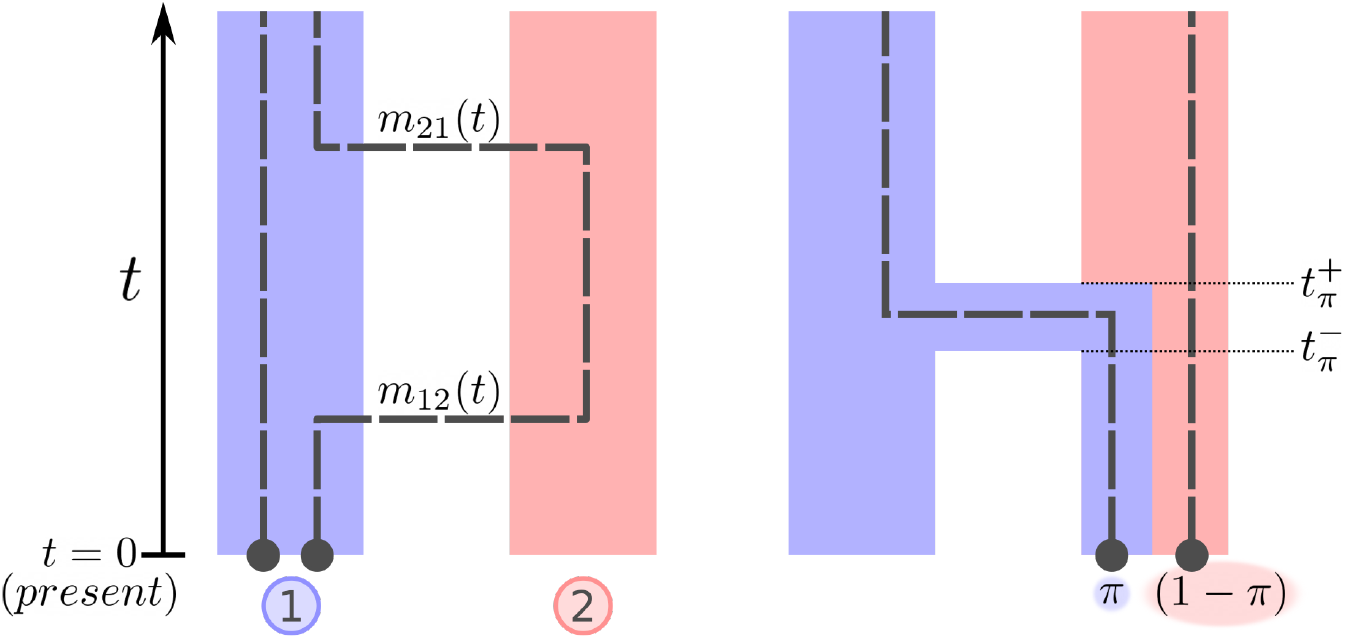
Notations for continuous migration (on the *left*) and pulse migration (on the *right*) models. In the continuous migration case, *m_ij_* is the migration rate from population *i* to population j, backwards in time. In the pulse migration case, *π* is the probability of migration of a lineage, *t_π_* is the instantaneous time of migration, 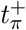 and 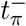 are the times right before and right after the migration pulse.

We here develop this concept for the case of two parental populations with admixture (continuous or pulse). We denote the two parental populations by *S*_1_(*t*) and *S*_2_(*t*). At any time *t*, a lineage ancestral to the observed population *S_m_* is either in population *S*_1_(*t*) or in population *S*_2_(*t*), due to migration. Within populations *S*_1_(*t*) and *S*_2_(*t*) lineages are fully exchangeable, which means that every pair of lineages from the same population has the same probability of coalescence. Effective population sizes of *S*_1_(*t*) and *S*_2_(*t*) are *N*_*L*1_(*t*) and *N*_*L*2_(*t*) respectively. *N*_*L*1_(*t*) and *N*_*L*2_(*t*) are what we define as *local effective population sizes*, i.e they represent the effective number of individuals in the populations at time *t* after discounting for the effect of migration. In other words, this is the rate of coalescence of two lineages conditional on both of them being in the given population.

If two lineages are in the same population *S_i_*(*t*) (*i* = 1, 2) at time *t*, they can coalesce with rate 1/*N_Li_*(*t*). If they are in different populations, coalescence is not possible between them. Conditional on two lineages having not coalesced by time *t*, let *P*_1_(*t*) and *P*_2_(*t*) be the probabilities that two lineages are in the population *S*_1_ and population *S*_2_ respectively. Let *P*_0_(*t*) be the probability that the two lineages are in different populations. Then the coalescence rate between a pair of lineages at time *t* is

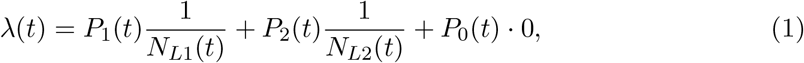

and the *ordinary effective population size* is

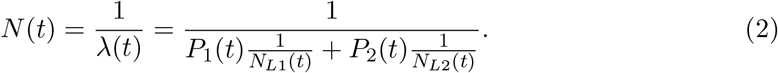

The condition that the sampled population *S_m_* is *S*_1_(0), is equivalent to setting the initial conditions of probabilities *P_i_* (*i* = 0, 1, 2) to

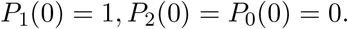

The dependence of *P_i_* on the time of observation is natural, because probabilities of migration might change over time, and even if they are constant, the cumulative amount of migration changes over time.

So, as shown above there is a clear difference between the *local* effective population size (*N*_*L*1_(*t*) and *N*_*L*2_(*t*)) of parental populations and the *ordinary* effective population size of an observed admixed population (*N*(*t*)). The estimates of effective population size obtained by PSMC and similar methods are estimates of the ordinary effective population size (*N*(*t*)) and not local effective population size (*N*_L_(*t*)).

### 2.2 Continuous and pulse migration

Equation 2 defining the ordinary effective population size, depends on the probabilities *P_i_*(*t*) (*i* = 1, 2, 0) of two lineages being in population *i* at time *t*, given they have not coalesced at *t*. These probabilities can be found by solving a set of differential equations. The past dynamics of two lineages is described by a coalescent model, which is a Markovian process going back in time. There are four possible states for this process:

- both lineages are in the first population at time *t* with probability *p*_1_(*t*),
- both lineages are in the second populations at time *t* with probability *p*_2_(*t*),
- the lineages are in different populations at time *t* with probability *p*_0_(*t*),
- the lineages have coalesced by time *t* with probability *p_c_*(*t*). This is an absorbing state.

Transitions between the first three states are possible through migration (either continuous or pulse). Transitions into the last absorbing state occur through coalescences, and *p_c_*(*t*) = 1 – *p*_1_(*t*) – *p*_2_(*t*) – *p*_0_(*t*). By definition of conditional probabilities,

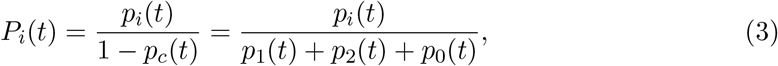

for *i* = 1, 2, 0.

The continuous migration rate *m_ij_*(*t*) (for *i* = 1, *j* = 2 or *i* = 2, *j* = 1) is the rate with which a single lineage from population *S_i_*(*t*) moves into population *S_j_* (*t*), backwards in time. Considered forward in time, these migration rates correspond to the fraction of population *S_i_* made of lineages from population *S_j_* (*i* ≠ *j*), when scaled in units of a reference effective population size *N*_0_ >> 1. This is the same definition as used in standard coalescence models with migration [Slatkin, 1982, 1987, Notohara, 1990, Wilkinson-Herbots, 1998] including Hudson’s ms simulator [Hudson, 2002].

Henceforth, we omit the dependence of these functions on *t* in our notations to improve readability. From standard definitions of the coalescent with migration, we then have the following system of differential equations:

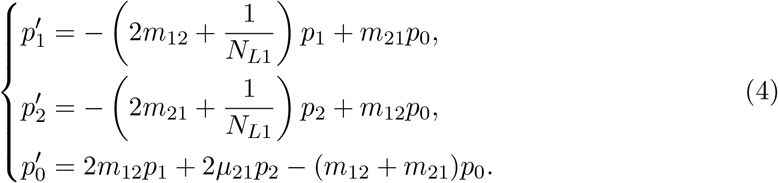

where 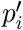 indicates the derivative of *p_i_* with respect to *t*.

Pulse migration acts instantaneously at time *t_π_*. Backwards in time, it drags a lineage from one population to the other with a certain probability *π*. Forward in time, *π* is the proportion of a recipient population made up of individuals from the donor population due to pulse admixture. Assume that the donor population is population 1 and the recipient population is population 2. We write 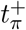 to indicate the time right before the pulse migration and 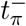 to indicate the time right after the pulse migration, forward in time (see Figure 1 for clarification). Then the probabilities *p_i_* (*i* = 1, 2, 0) change as follows

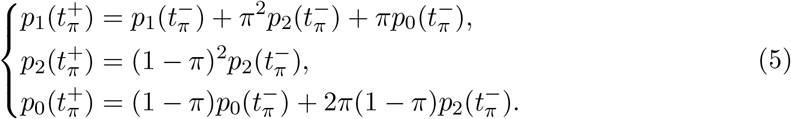

The parameter *π* is equivalent to the parameter 1 – *p* of the -es switch in Hudson’s ms simulator [Hudson, 2002].

### 2.3 Disentangling the effect of migration on effective population size

Assume that we observe two populations 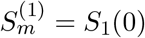 and 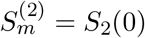, which had ancestral admixture with each other. Writing equation 2 for samples from both populations, we get the system of equations relating the *ordinary effective population size* of 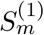 and 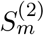 (*N*_1_ and *N*_2_) with the *local effective population size* of each of the two parental populations (*N*_*L*1_ and *N*_*L*2_).

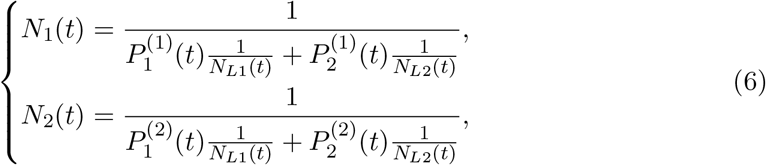

where 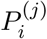 is the probability that both ancestral lineages from population 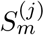 are in population *i* (see equation 3). These functions can be derived from equation 2 by setting initial conditions to 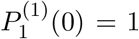 for the first populations and 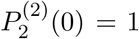 for the second population.

As we already mentioned, PSMC or similar methods can be used to estimate the ordinary effective population size, *N*(*t*). Given samples from two admixed populations, the underlying local effective population size (*N*_*L*1_ and *N*_*L*2_) of their parental populations *S*_1_ (*t*) and *S*_2_(*t*) can be estimated from equation 6.

Unfortunately, there is no closed form solution of equations 4 and 6. PSMC approximates historical effective population size with a piece-wise constant function. In our inference, we assume that migration rates are constant in each time interval, and similarly to PSMC, we approximate local effective population sizes with a piece-wise constant trajectory. So, instead of solving equation 6, we calculate piece-wise constant functions *N*_*L*1_(*t*) and *N*_*L*1_(*t*) such that the probabilities to coalesce within each time interval is the same as inferred by PSMC. In more details, for two lineages from population *i* the probability of coalescence 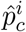 within [*t*_1_, *t*_2_] inferred by PSMC is

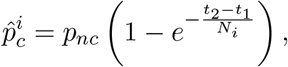

where *p_nc_* is the probability that two lineages have not coalesced by time *t*_1_.

From equation 4 the probability 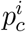 that two lineages from population *i* coalesce within the interval [*t*_1_, *t*_2_] is

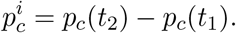

And we fit *N*_*L*1_ and *N*_*L*2_ so that

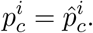

### 2.4 Estimating migration rates and split time

In the previous subsection we show how one can calculate local effective population sizes for given values of migration rates and split time. Of course, it is also desirable to estimate these parameters, because they are often unknown. Our method fits the joint site frequency spectrum (SFS) of two diploid individuals representing two populations. Let *f_i,j_* (*i, j* = 0, 1, 2) be the probability that a variable site has *i* derived alleles in the first individual and *j* derived alleles in the second individual. Notice that *f*_0,0_ and *f*_2,2_ are excluded because they correspond to non-variable sites. Then the probabilities *f_i,j_* define a multinomial distribution.

Let **n** = {*n*_0,1_, *n*_1,0_, *n*_1,1_, *n*_1,2_, *n*_2,1_} be the site frequency spectrum from the data, i.e. *n_i,j_* is the non-normalised counts of sites with *i* and *j* derived alleles in the first and second individual, respectively. We consider the composite likelihood function (which ignores possible correlations between the sites) given by the multinomial distribution:

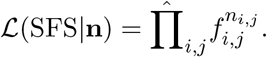

The theoretical SFS for a given set of parameters and PSMC trajectories can be computed by numerically solving a set of linear differential equations describing a Markov process with 44 states. These states describe all possible ways in which lineages of a coalescent tree with four tips (two samples from each of two populations) can be distributed among populations. These states include the possibility of coalescence between lineages and migration between populations through time (details are given in the Appendix A).

### 2.5 Software implementation

Our method for estimating underlying local effective population size is implemented in Python 3 under the name MiSTI. The implementation is available at https://github.com/vlshchur/MiSTI and distributed under GNU GPL3.

## 3 Results

### 3.1 Obtaining local effective population size from the ordinary effective population size estimated by PSMC

In this section we demonstrate the effect of migration on effective population sizes, and we will qualitatively assess the PSMC inference of historical effective population size trajectories.

We used ms [Hudson, 2002] to simulate two population size trajectories: one trajectory (population 1) has constant size through time after the population split, and the second trajectory (population 2) has a bottleneck after the split, followed by a recent population expansion. Using these population size trajectories, we simulated symmetric migration between populations, as well as unidirectional migration. For each simulation, we show: 1) the ordinary effective population size of both populations (*N*_1_ and *N*_2_) calculated using Equation 6, 2) the ordinary effective population size estimated using PSMC (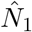 and 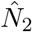), and 3) the local effective population size (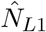 and 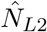) obtained with MiSTI by correcting the effective population size trajectories for the effect of migration.

Continuous, bidirectional migration between populations 1 and 2 from the present until the split time generally increases the ordinary effective population size (*N*_1_ and *N*_2_) relative to the simulated local population size (*N*_*L*1_ and *N*_*L*2_, Figure 2A). However, notice that population 1 has a decreased effective population size relative to its simulated local size, during the population 2 bottleneck (Figure 2A). This decrease in genetic variation observed in population 1 is caused by the possibility that lineages from population 1 go through the bottleneck in population 2, where coalescence rates are increased.

**Figure 2:**
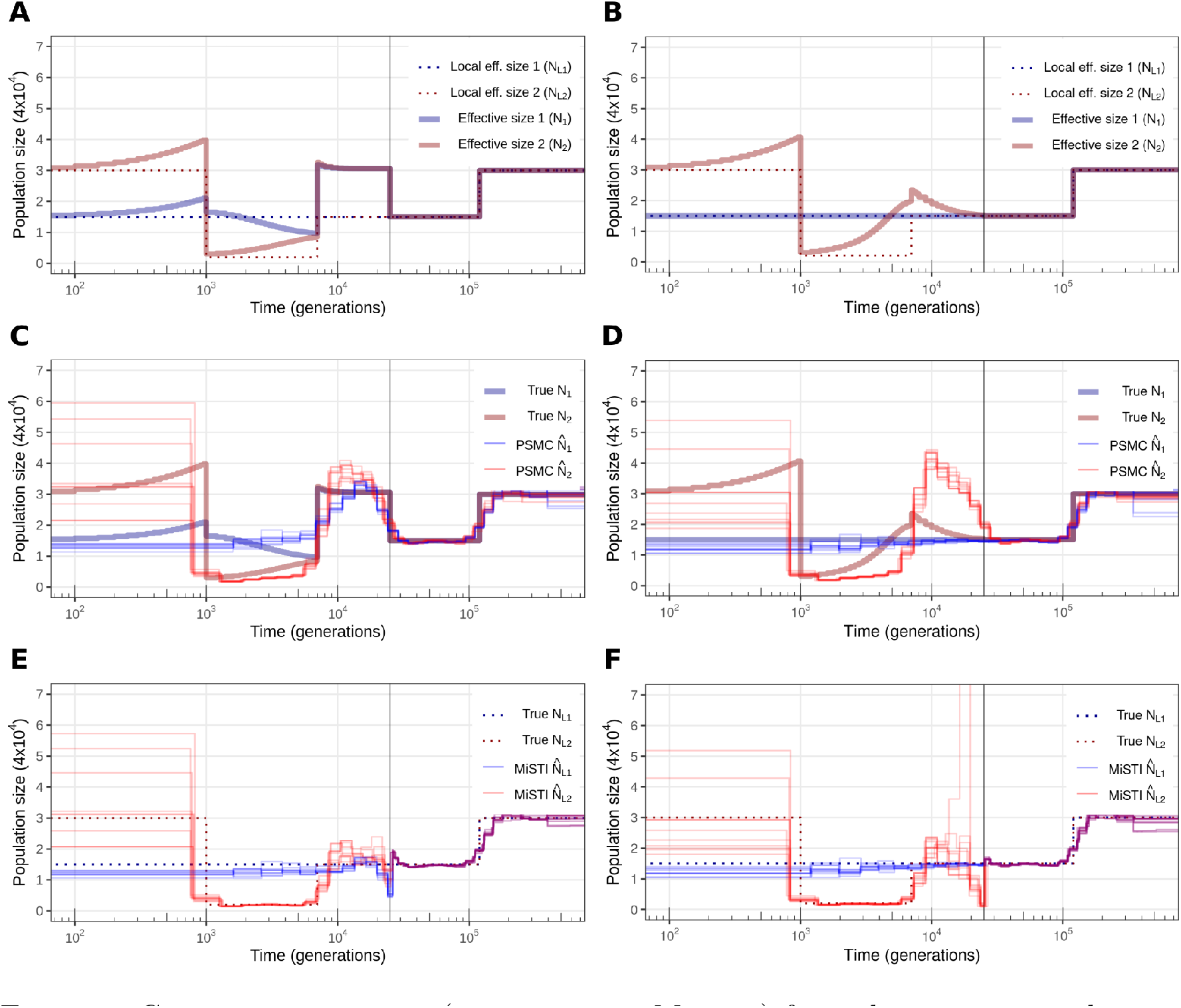
Continuous migration (ms parameter *M_ij_* = 2) from the present to split time (25000 generations, indicated by vertical bar). (A,C,E) Bidirectional migration. (B,D,F) Uni-directional migration from population 1 to population 2. (A,B) Simulated local effective population sizes and ordinary effective population sizes calculated according to equation 6. (C,D) True ordinary effective population size from A,B, and estimated by PSMC. (E,F) True local population sizes from A,B, and estimated by MiSTI.

In this scenario, PSMC generally estimates the ordinary effective population size (Figure 2C) well, despite the smoothing of instantaneous populations size changes that has been described previously [Li and Durbin, 2011]. We also note that in Figure 2A, the historical effective population size of populations 1 and 2 coincide before the bottleneck, but PSMC trajectories do not coincide (Figure 2C). We discuss this possible bias in PSMC below.

Continuous, unidirectional migration from population 1 to 2, generates an increase in the ordinary effective population size of population 2 (Figure 2B), which is detected by PSMC (Figure 2D). In this scenario, the inferred PSMC trajectory underestimates effective population size of population 2 (the one receiving migrants) during the bottleneck, and overestimates it before the bottleneck. We hypothesize that this effect, as well as the discordance of PSMC curves prior to the bottleneck in Figure 2C, could be due to the violation of the SMC model assumption that samples come from a single panmictic population, as we explore further in the next section.

Applying MiSTI correction of PSMC curves with the known migration rates and split times used in the simulations, to estimate local population sizes (*N*_*L*1_ and *N*_*L*2_), recovers trajectories similar to the simulated ones (Figures 2E,F).

Pulse migration also increases historical effective population size. A single pulse of migration at time zero will cause effective population size to increase monotonically backwards in time until the time when populations split (Figure 3A,B). Similar to the continuous migration case discussed above, PSMC detects this increase in historical effective population size, although it underestimates the extreme peak of effective population size preceding the population split (Figure 3C,D). This underestimation is due to an smoothing effect of the PSMC method, which has been described [Li and Durbin, 2011]. MiSTI recovers the local effective sizes of populations 1 and 2, slightly underestimating it when the PSMC smoothing underestimated the peak in effective population size (Figure 3E,F).

**Figure 3:**
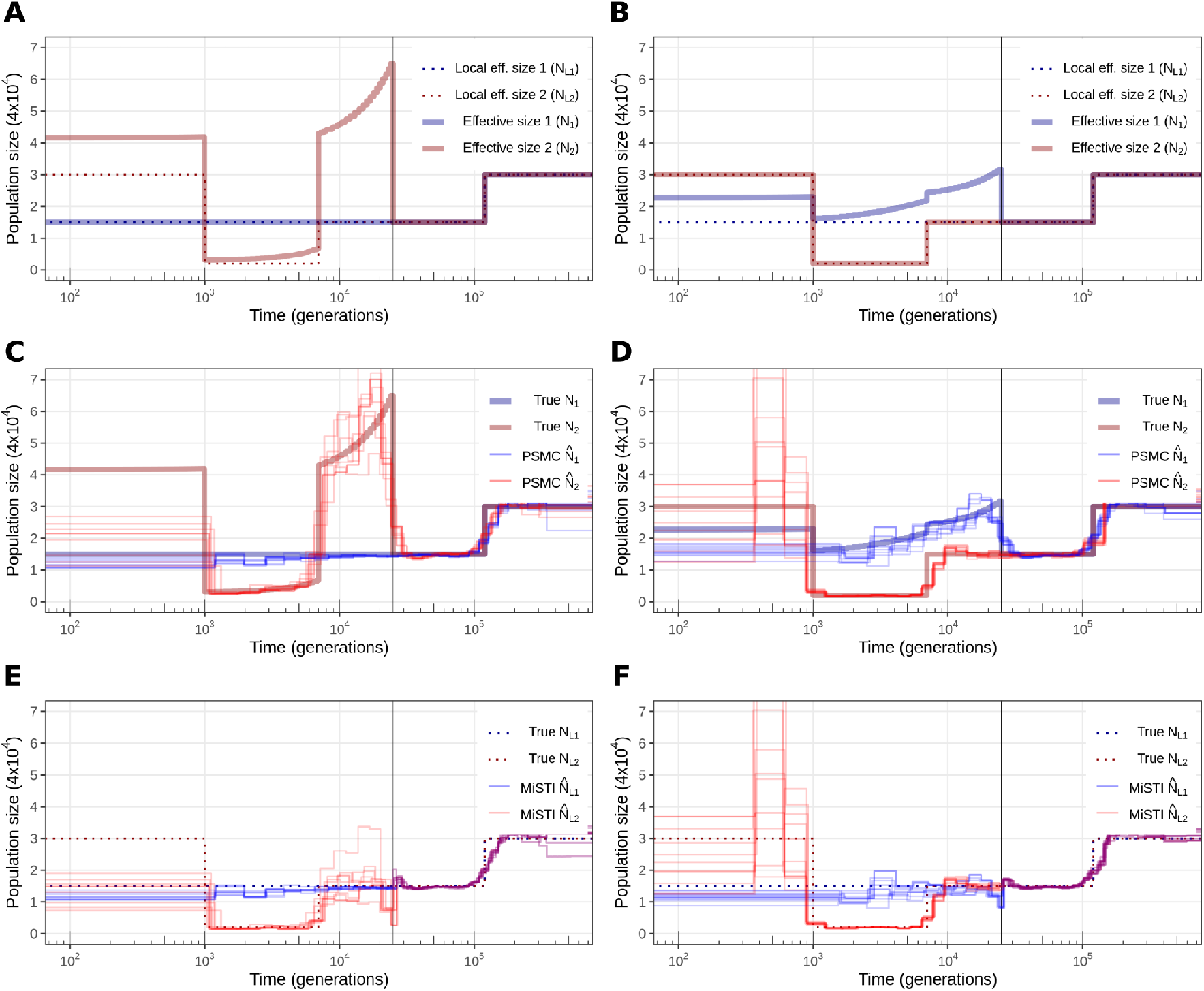
Pulse migration of 20% at the present time. (A,C,E) pulse from population 1 to population 2. (B,D,F) pulse from population 2 to population 1, forward in time. (A,B) Simulated local effective population sizes and ordinary effective population sizes calculated according to equation 6. (C,D) True ordinary effective population size from A,B, and estimated by PSMC. (E,F) True local population sizes from A,B, and estimated by MiSTI.

### 3.2 Transition matrices and the assumption of a single panmictic population in PSMC

PSMC assumes a model of a single panmictic population. When data comes from a structured population, PSMC can often find a best fitting transition matrix that has a stationary distribution equivalent to that of a single population with the same effective population size [Chikhi et al., 2018]. However, in the previous section, we showed one example where the ordinary effective population size inferred by PSMC was strongly biased (“population 2” in Figure 2D). This reveals that the stationary distribution of the transition matrix fitted by the HMM underlying PSMC is different from the true distribution.

To investigate this bias in PSMC, we simulated a single panmictic population with the same effective population size as population 2 (*i.e*. following the trajectory of *N*_2_ in Figure 2D). In other words, this population has the same stationary distribution of the coalescence time transition matrix as population 2, but it did not receive any migrants. Let us call this population *P*, for panmictic. We found that the empirical transition matrix of population *P* (Figure 4C) differs more from the empirical matrix of population 2 (Figure 4E) than the transition matrix inferred by PSMC (Figure 4D). This indicates that the matrix inferred by PSMC is a better fit of the empirical matrix, and therefore the PSMC bias we detected is not due to an optimization problem.

**Figure 4:**
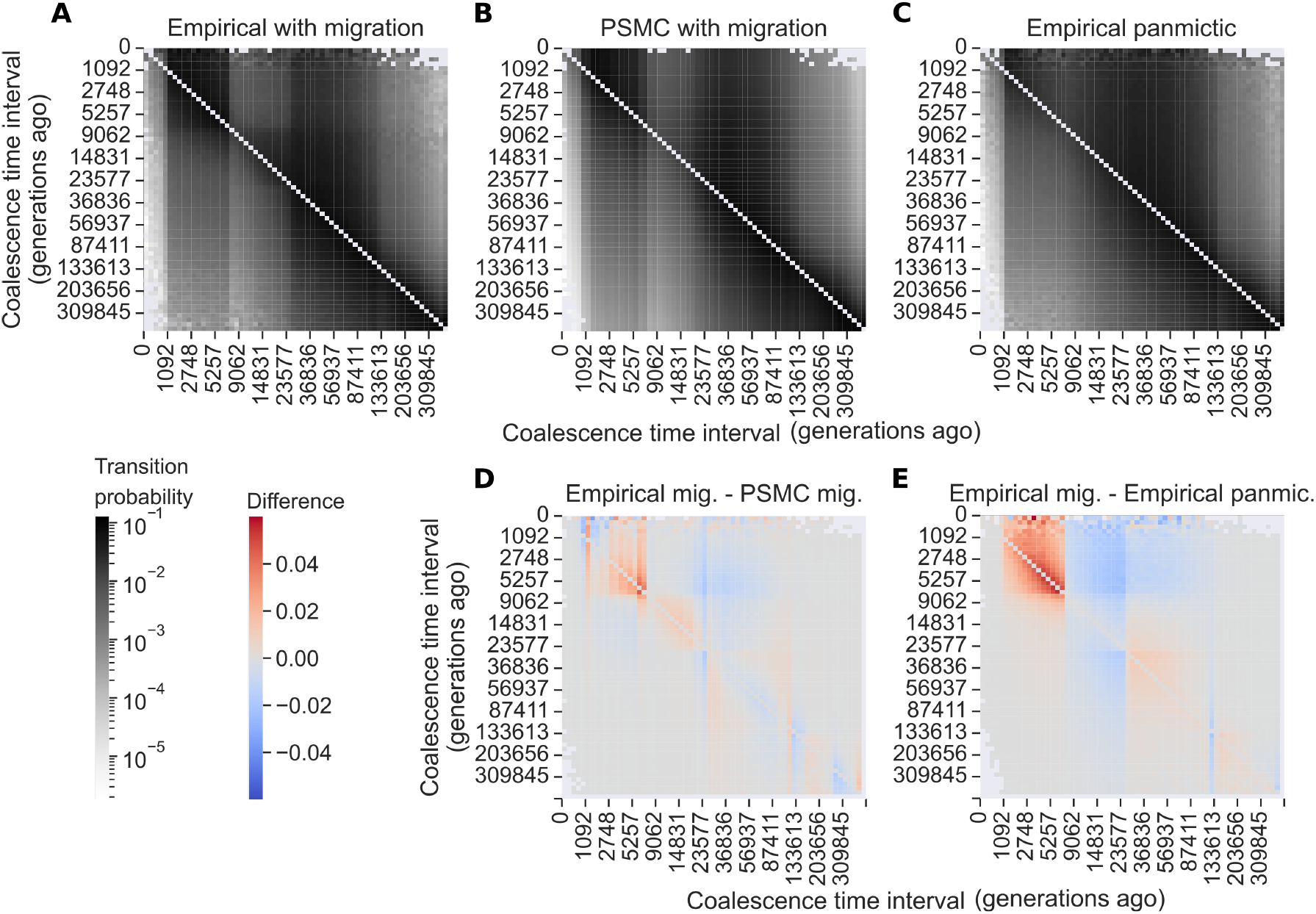
Matrices representing the (empirical or estimated) probability of transitions from one coalescence time to another along a sequence. (A) Transition matrix from simulated data from a population receiving migrants (population two of Figure 2B). (B) Transition matrix estimated by PSMC. (C) Transition matrix from simulated data from a single population (no migration) with historical effective population size equal to population two of Figure 2B. (D) Difference between matrices A and B. (E) Difference between matrices A and C.

Next, we compare the empirical transition matrix from population 2 (Figure 4A) to the transition matrix inferred by PSMC (Figure 4B). One difference between those matrices is that the PSMC matrix (Figure 4B) shows mostly vertical bands, while the true transition matrix (Figure 4A) shows a horizontal band corresponding to the time between the bottleneck and the split of populations (7-25k generations).

The horizontal band in the true transition matrix in Figure 4A is caused by a correlation in coalescence times between adjacent sites that can not arise in a panmictic model, assumed by PSMC. If two lineages coalescence during the bottleneck at a site, then there is an increased probability that both these lineages are in the bottlenecked population at other sites. Furthermore, if recombination happens during the bottleneck period, and both lineages are in the bottlenecked population, then there is an increased probability that the two lineages again will coalesce in this time interval in the next site after recombination. This contrasts with a standard SMC coalescence model in a panmictic population, in which the time of coalescence in site *i* + 1 is independent of the coalescence time in site *i*, conditional on it being older than the time of recombination between the sites. A structured model with a bottleneck, therefore, creates a correlation structure that cannot be modeled by the standard SMC model used in PSMC.

The bias in PSMC can therefore be explained by the fact that PSMC fits the transition matrix, and not its stationary distribution. Importantly, it fits the transition matrix of a panmictic model. As we mentioned previously, in many cases, fitting the best panmictic transition matrix also fits the best stationary distribution, but in this case, the best fit of a panmictic transition matrix by PSMC leads to a very different stationary distribution.

We also note that the single population model assumed by PSMC differs from the structured model with migration in the way that recombination rate scales with effective population size through time. In a single population model, an increase in effective population size (*N*) increases the effective recombination rate (*ρ* = 4*Nr*, where *r* is the rate of recombination per locus per generation). In a model with two populations, an increase in effective population size that is due to migration would decrease the effective recombination rate, since two lineages can only recombine when they are in the same population. In other words, in a model with a single population, the effective recombination rate scales with the effective population size, while it is not necessarily true in a model with two populations and migration, where it can in fact scale inversely with effective population size.

### 3.3 Correcting European effective population size for the Neanderthal component

In this section, we show an application of the MiSTI correction of PSMC curves using known split time and admixture rates.

Non-African human populations admixed with Neanderthals 52-58 thousand years ago [Prüfer et al., 2014]. Villanea and Schraiber [2019] recently reported that it is likely that there were multiple admixture events, but for simplicity we consider the case of a single pulse admixture event. This admixture would result in a number of very old coalescences which would increase the overall estimate of effective population size. In order to estimate the local effective population sizes of non-African populations, we need to correct for this admixture. One way of doing that is to call and mask the Neanderthal introgressed regions in a modern genome before running PSMC. Alternatively, one can estimate the proportion of Neanderthal ancestry in a modern genome, and use MiSTI to correct its PSMC trajectory for that admixture proportion. We compare these two approaches to verify that MiSTI’s correction of effective population size is consistent with masking of known tracts of introgression.

We removed the Neanderthal tracts in an European genome (CEU population) and confirmed that the PSMC trajectory inferred from the Neanderthal-masked genome has lower effective population sizes than the non-masked, original genome (Figure 5). Next, we used MiSTI to correct the PSMC trajectories of the non-masked European genomes assuming 1.5% Neanderthal introgression, which has been reported by Steinrücken et al. [2018], and 3.0%, which was the previously reported estimate [Green et al., 2010] (Figure 5). Correcting the CEU PSMC trajectory for 1.5% Neanderthal admixture using MiSTI gives very similar estimates of local effective population size as masking the known regions of Neanderthal ancestry from that same genome. This suggests that MiSTI, at least in this case, correctly recovers the effective population size of parental populations, when applied to real data.

**Figure 5:**
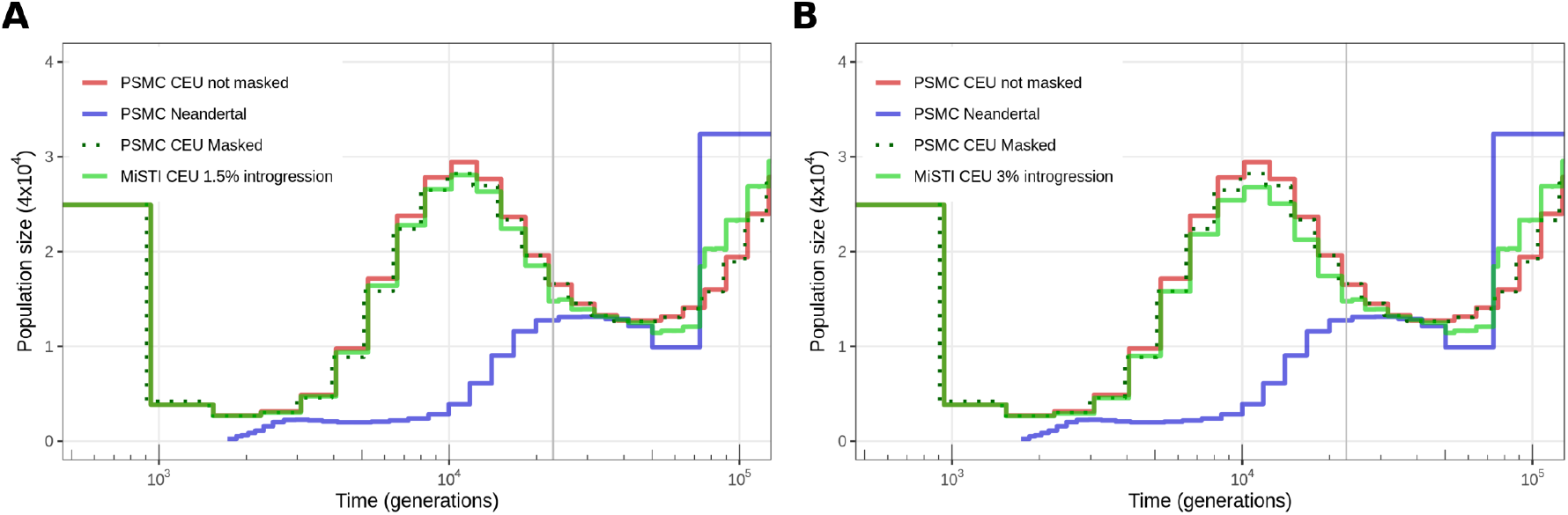
(A) MiSTI correction of PSMC effective population size trajectory of a genome from the CEU population assuming 1.5% introgression and (B) 3% introgression from Neanderthals.

In the Appendix B, we show another application of MiSTI to obtain local effective population size by correcting PSMC curves for the effect of migration, using known parameters from *Puma concolor* populations, and we discuss limitations of applying MiSTI to that case.

### 3.4 Estimating split time in human-like simulations

Most often, split times and migration rates are unknown, and MiSTI can be used to estimate these parameters from PSMC curves combined with a joint (2D) site frequency spectrum for the pair of samples. In this section, we estimate split times from simulations replicating effective population size trajectories similar to those of human populations.

We simulated populations approximating the historic effective population size of human populations (Dinka, San, Sardinian, French and Han) (see “Simulations” in Appendix A). Briefly, we simulated historic effective population size similar to the estimated by PSMC for each of those populations (Figures 6A, 7A, 8A). We simulated population splits at various times, with no migration following the split. We then estimate these split times using MiSTI and the TT-method [Sjödin et al., 2021] (Figures 6B, 7B, 8B). We found that the TT method estimates negative split time in simulations where the split time happens during or immediately at the end of the bottleneck (Figure 8B), as has been previously described [Sjödin et al., 2021]. In other scenarios of intermediate split times, the TT method largely overestimates the split times, due to violations of the assumption of constant effective population sizes in the ancestral population [Sjödin et al., 2021]. In contrast, MiSTI provides substantially less biased estimates.

**Figure 6:**
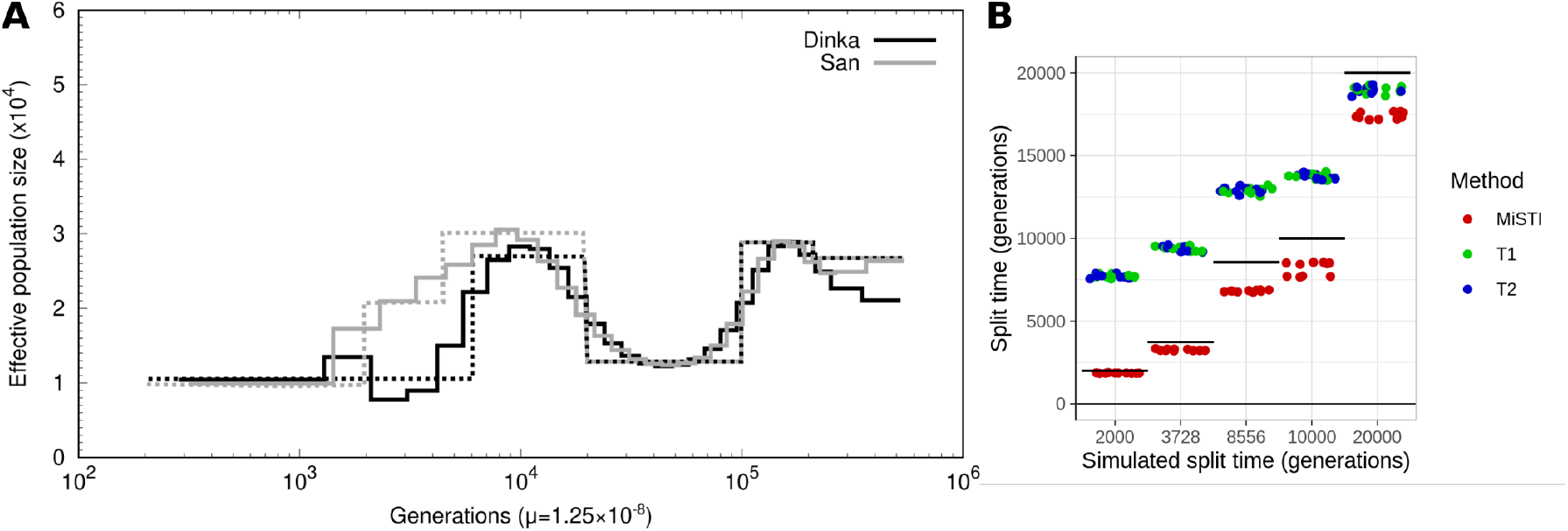
Simulations of San-Dinka split, no migration. (A) Continuous lines show effective population size inferred by PSMC from real data, dotted lines show simulated population sizes that approximate the inferred trajectory. (B) Inferences from MiSTI and the TT method for ten replicate simulations of each split time. 3728 generations was the split time inferred by MiSTI from real data; 8556 was the split time inferred by the TT method (see Table 2) - note that when we simulate 3728 generations, the TT method infers close to 8556 generations.

**Figure 7:**
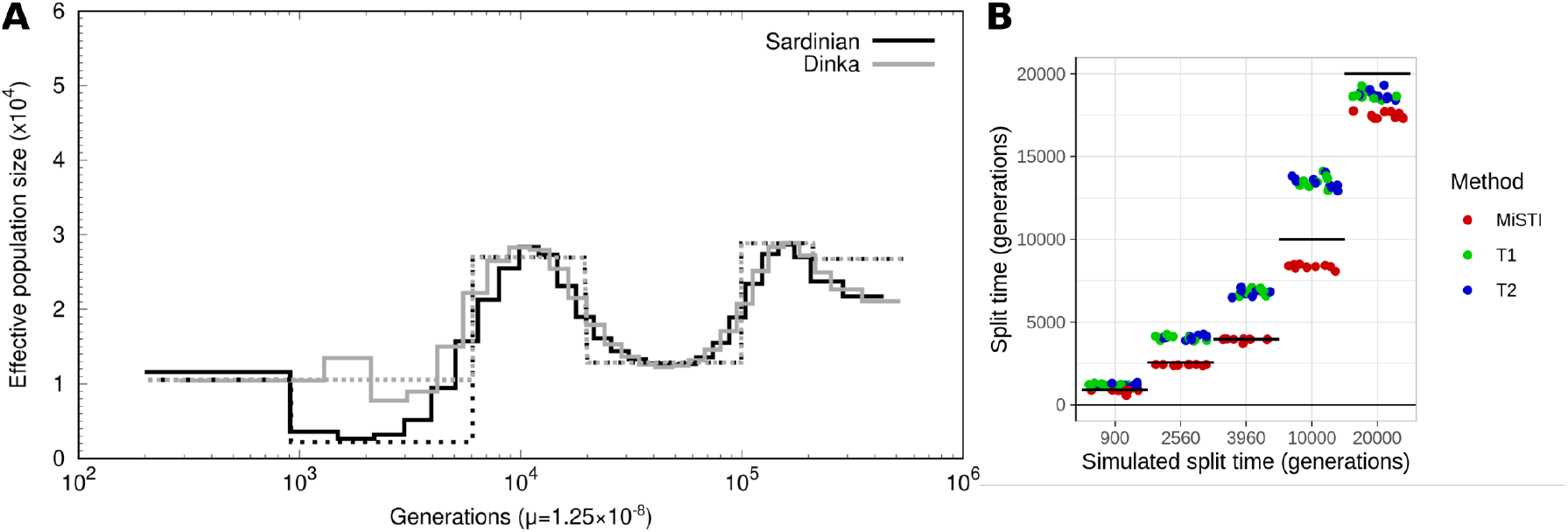
Simulations of Dinka-Sardinian split, no migration. (A) Continuous lines show effective population size inferred by PSMC from real data, dotted lines show simulated population sizes that approximate the inferred trajectory. (B) Inferences from MiSTI and the TT method for ten replicate simulations of each split time. 3960 generations was the split time inferred by MiSTI from real data; 2560 was the split time inferred by the TT method (see Table 3).

**Figure 8:**
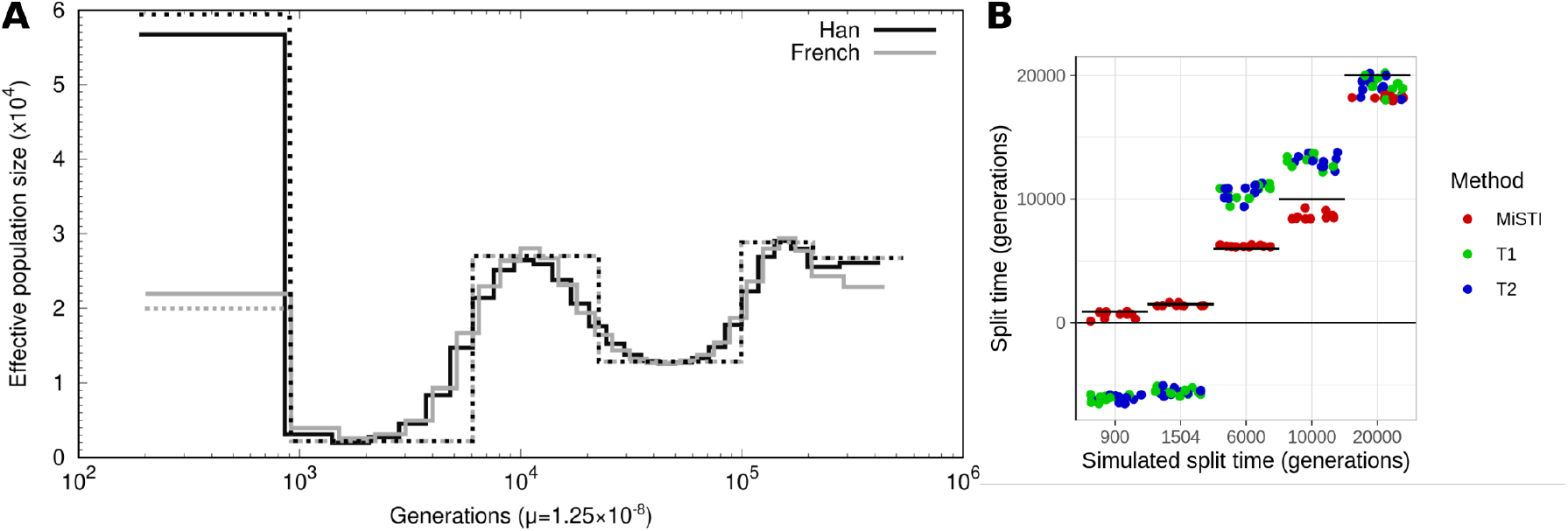
Simulations of Han-French split, no migration. (A) Continuous lines show effective population size inferred by PSMC from real data, dotted lines show simulated population sizes that approximate the inferred trajectory. (B) Inferences from MiSTI and the TT method for ten replicate simulations of each split time. 1504 generations was the split time inferred by MiSTI from real data; −3566 was the split time inferred by the TT method (see Table 1).

### 3.5 Estimating split time and migration rates from human data

Here, we estimate split times and migration rates from real data from the same populations we simulated in the previous section. We used MiSTI to estimate split-times and migration rates between the Han Chinese and French (Table 1), Dinka and Sardinian (Table 3), San and Dinka (Table 2) and San and Sardinian (Table 4) populations. From MiSTI, we recorded the maximum composite likelihood values of three models: no migration, unidirectional migration in each direction, and bidirectional migration. In all models with migration, we assumed a constant rate of migration between the split time and the present.

**Table 1:** MiSTI estimates of split times and migration rates between the Han Chinese and French populations in models with bidirectional migration (top row), unidirectional migration, or no migration (bottom row).

| m1<br>French to Han | MiSTI |  |  | TT<br>split time<br>(generations) |
| --- | --- | --- | --- | --- |
|  | m2<br>Han to French | split time<br>(generations) | log(lik) |  |
| $8.37 \times 10^{-9}$ | 2.92 | 1505 | -2331 | - |
| - | 2.92 | 1505 | -2331 | - |
| 3.84 | - | 1505 | -2373 | - |
| - | - | 1505 | -2787 | T1 = -3587<br>T2 = -3545 |

**Table 2:** MiSTI estimates of split times and migration rates between the San and Dinka populations in models with bidirectional migration (top row), unidirectional migration, or no migration (bottom row).

| m1<br>Dinka to San | MiSTI |  |  | TT<br>split time<br>(generations) |
| --- | --- | --- | --- | --- |
|  | m2<br>San to Dinka | split time<br>(generations) | log(lik) |  |
| 2.5 | $2.03 \times 10^{-9}$ | 3729 | -4381 | - |
| 2.5 | - | 3729 | -4381 | - |
| - | 1.49 | 3210 | -4582 | - |
| - | - | 3001 | -4607 | T1 = 8582<br>T2 = 8527 |

**Table 3:** MiSTI estimates of split times and migration rates between the Dinka and Sardinian populations in models with bidirectional migration (top row), unidirectional migration, or no migration (bottom row).

| m1<br>Sardinian to Dinka | MiSTI |  | split time<br>(generations) | log(lik) | TT |
| --- | --- | --- | --- | --- | --- |
|  | m2<br>Dinka to Sardinian |  |  |  | split time<br>(generations) |
| 2.63 | 9.96 |  | 3963 | -5337 | - |
| 6.92 | - |  | 3484 | -5819 | - |
| - | 14.40 |  | 2286 | -10877 | - |
| - | - |  | 2264 | -13166 | T1 = 2529<br>T2 = 2577 |

**Table 4:**
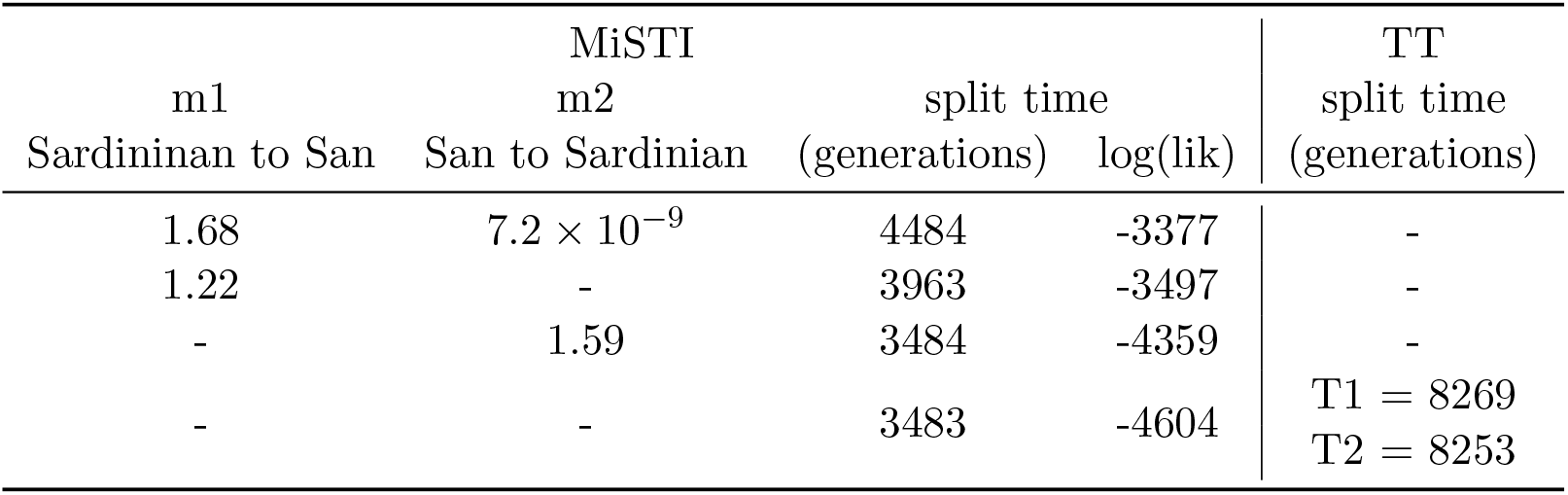
MiSTI estimates of split times and migration rates between the San and Sardinian populations in models with bidirectional migration (top row), unidirectional migration, or no migration (bottom row).

For Han-French divergence, the model with the highest composite likelihood was one with a split time of 1505 generations (*i.e*. 43,645 years ago assuming 29 years per generation) and a mostly unidirectional migration rate of 2.92 from Han to French (Table 1). We also replicate the results from Sjödin et al. [2021], in which the TT method infers nonsensical negative split times between Han and French. The unidirectional migration inferred from Han to French is in line with current models of the peopling of Europe through waves of farmers coming from central Eurasia [Haak et al., 2015].

The best fit model for the San-Dinka population pair includes a split time of 3729 generations ago (*i.e*. 108,141 years ago assuming 29 years per generation), and mostly unidirectional migration from Dinka to San. For the same data, the TT method infers a much larger split time (over 8500 generations ago) (Table 2, see also Appendix C for a validation of this result with simulations).

The Dinka-Sardinian split time inferred by MiSTI is approx. 3963 generations ago, with bidirectional migration between these populations. The migration rate detected from Dinka to Sardinian is in line with previous results indicating migration from sub-Saharan Africa to South Europe [Moorjani et al., 2011]. In this case, in contrast to the previous case, the TT method infers a more recent split time than MiSTI (2550 generations ago, see Table 3). The split time between San and Sardinian is older (approx. 4484 generations ago, see Table 4). We notice that these estimates are not strictly compatible with a population tree, which likely is a consequence of complex ancestral population structure and migration between populations that is not modeled here, including archaic admixture into Sardinians. We note that archaic admixture will tend to inflate divergence time estimates, so the true divergence times might be smaller than our estimates, particularly for the splits between Sardinians and the two African populations.

## 4 Discussion

### 4.1 The MiSTI method

The coalescent effective population size, defined as the reciprocal of the coalescence rate, is proportional to the census population size in a panmictic model, but can be very different from it when there is migration. The idea of disentangling the effect of migration on effective population size has been explored before. For example, Wang and Whitlock [2003] introduced methods to jointly estimate the local effective population size and migration rates from samples taken over time and space. Here, we are motivated by the same idea of disentangling migration and effective population size, and we do so in the context of inferring changes in effective population size through time from present-day samples of different populations, with methods such as PSMC [Li and Durbin, 2011].

We defined the ordinary effective population size of an admixed population as a function of the local effective population size of its parental populations. The local effective population size corresponds to the effective population size of unadmixed individuals from the parental populations. We developed a method, MiSTI, that uses the ordinary effective population sizes (*e.g*. estimated by PSMC [Li and Durbin, 2011]) of two samples from different populations that exchanged migrants, together with their joint SFS, to estimate the local effective population sizes and migration rates.

We note that MiSTI depends on the results of PSMC, and will be subject to its biases. MiSTI relies on a model of a population split possibly followed by migration, which itself violates the assumption of panmixia made in PSMC and similar methods. We have shown that PSMC estimates are particularly sensitive to this violation when there is asymmetric migration (Figure 2D). In other scenarios, us (Figures 2C, 3C, D) and others [Chikhi et al., 2018] have shown that PSMC provides good estimates of historical effective population size in the presence of population structure. Other studies have shown, using simulations, that estimates from PSMC and other methods can be biased even in the absence of population structure [Spence et al., 2018]. We have not extensively explored those biases here, but we note that as less biased methods are developed, MiSTI can be adapted to use those and thus improve its inference of local effective population size, split times and migration rates.

When applied to infer population split times and migration rates in human populations, MiSTI helps settle a previous controversy. Schlebusch et al. [2017] found surprisingly deep divergence times for some Southern African populations, including the San. Their estimate for the split time between Dinka and San is 255 ± 5 years ago (Figure 3C in Schlebusch et al. [2017]). We applied the TT method used in Schlebusch et al. [2017] to estimate the San-Dinka split time in our data and we found a similar result (split time 8554 generations ago, or 248 thousand years with 29 years per generation, Table 2), replicating their results. However, with MiSTI, we estimate a much more recent split time around 3729 generations ago (108 thousand years ago with 29 years per generation, Table 2). Our estimate is similar to the estimates of the earliest population divergence among modern human populations obtained with methods such as MSMC [Pagani et al., 2016, Fan et al., 2019], and momi2 [Kamm et al., 2020] (see Bergström et al. [2021] Figure 2C for a synthesis of estimates from various studies).

The TT method makes a strong assumption that there are no changes in population size in the ancestral population, before the population split [Schlebusch et al., 2017, Sjödin et al., 2021]. We simulated historical effective population size similar to the one estimated by PSMC for San-Dinka, and showed that the TT method strongly overestimates split times in this scenario (Figure 6). Notably, when we simulated data assuming a split time of 3728 generations ago (as inferred by MiSTI), the TT method estimated a split time close to the one it estimated from the real data (8556 generations ago), showing that the previously reported deep split time estimated by the TT method is in fact likely an estimation artifact. The TT method can be highly biased because of the assumption of constant population size and should not be applied to populations that may have experienced changes in effective population size over time.

The application of MiSTI to human data also illustrates the importance of including migration a the model is used to infer split times. In all cases (Tables 1-4), composite likelihoods were higher in models that allowed migration, and a difference of 1000 or more generations is seen in some split times inferred with models that include migration (Tables 3-4). Inferring asymmetric migration is also an interesting feature of MiSTI aimed at determining the direction of gene-flow.

### 4.2 Why estimate local effective population size?

Finally, we would like to highlight the broader relevance of disentangling the effect of migration on effective population size. The ordinary effective population size is important for understanding patterns of neutral genetic variability. It is a good predictor of summary statistics of neutral genetic variation such as the expected heterozygosity and average number of pairwise differences. However, for questions related to the efficacy of selection or genetic drift, the effective population size defined in terms of the number of individuals and their variance in offspring number is what matters most, not the effective population size inflated by migration. Since the ordinary effective population size is generally increased by migration, recovering the local effective population size after accounting for the effect of migration will recover values that are more informative for selection dynamics and predictions regarding the efficacy of selection, such as the rate of purging of deleterious alleles.

Local effective population size is also often more relevant for conservation genetics than the ordinary effective population size, which reflects overall genetic diversity of a metapopulation. For example, a meta-population of an endangered species which occupies a fragmented habitat might have increased effective population size if considered as a whole. Their apparent high levels of neutral genetic diversity might be misleading regarding their fragile conservation status. If there is weak migration between isolated subpopulations, the ordinary effective population size of each subpopulation will be inflated by migration and will not be representative of the actual size of the local population. Correcting for the effect of population structure and migration provides measures of local effective population size that are closer to the effective number of breeding individuals in the population and thus more informative for conservation efforts.

## 5 Acknowledgements

VS worked on this paper within the framework of the HSE University Basic Research Program. AI was supported by the grant RFBR 20-29-01028. This material is based upon work supported by NIH grant R01GM138634 awarded to RN.

## A Supplementary Methods

### A.1 Markov process describing state of lineages in a coalescence tree with four tips

Let us encode a lineage state at time *t* by (*k, l, i*), where *k, l* ∈ {0, 1, 2}, *k* + *l* > 0 are the number of descendants of this lineage that were sampled (at time *t* = 0) from populations 1 and 2 respectively, and *i* ∈ {1, 2} is the population where the lineage is at time *t*. We do not consider equations corresponding to states with a single lineage, *i.e*. before the most recent common ancestor of the 4 samples ({2, 2, i}, *i* = 1, 2), because they do not contribute to variable sites.

Before split time there is one single ancestral population, so a similar approach holds but the last index *i* is not needed to encode a lineage. So, there are only 8 possible states of the Markov process, and an additional absorbing state (2, 2) which does not contribute to SFS. We proceed with the derivation of the case of two ancestral populations, as it is a more complex one.

Every Markov state {(*k_j_, l_j_, i_j_*)} is a set of lineages (enumerated with the index *j*, 1 < *j* ≤ 4) with the condition ∑*_j_ k_j_* = ∑*_j_ l_j_* = 2. At the time of observation *t* = 0 the initial state is {(1, 0, 1), (1, 0, 1), (0, 1, 2), (0, 1, 2)}. Two lineages (*k*_1_, *l*_1_, *i*_1_) and (*k*_2_, *l*_2_, *i*_2_) can coalesce only if *i*_1_ = *i*_2_, and the resulting lineage is (*k*_1_ + *k*_2_, *l*_1_ + *l*_2_, *i*_1_).

Let us consider the state *L* = {(1, 1, 1), (1, 0, 1), (0, 1, 2)}, and write the equation for the derivative *P_L_*(*t*) which is the change in the probability of the Markov process being in state *L* at time *t*.

Transitions into state L are possible from the following four states

- *L*_1_ = {(1, 0, 1), (1, 0, 1), (0, 1, 1), (0, 1, 2)} through coalescence of any of two lineages (1, 0, 1) and the lineage (0, 1, 1) with the total rate of coalescence 2/*N*_*L*1_,
- *L*_2_ = {(1, 1, 2), (1, 0, 1), (0, 1, 2)} through migration of the lineage (1, 1, 2) from population 2 into population 1 with the migration rate *m*_21_,
- *L*_3_ = {(1, 1, 1), (1, 0, 2), (0, 1, 2)} through migration of the lineage (1, 0, 2) with the rate *m*_21_,
- *L*_4_ = {(1, 1, 1), (1, 0, 1), (0, 1, 1)} through migration of the lineage (0, 1, 1) with the rate *m*_12_.

Transitions from state L are possible into four states

- *L*_5_ = {(2, 1, 1), (0, 1, 2)} through coalescence of the lineages (1, 1, 1) and (1, 0, 1) with the coalescence rate 1/*N*_*L*1_,
- *L*_2_ = {(1, 1, 2), (1, 0, 1), (0, 1, 2)} through migration of the lineage (1, 1, 1) from population 1 into population 2 with the migration rate *m*_12_,
- *L*_3_ = {(1, 1, 1), (1, 0, 2), (0, 1, 2)} through migration of the lineage (1, 0, 1) with the rate *m*_12_,
- *L*_4_ = {(1, 1, 1), (1, 0, 1), (0, 1, 1)} through migration of the lineage (0, 1, 2) with the rate *m*_21_.

So, the corresponding equation is

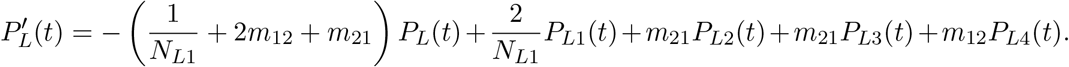

Mutation on a lineage (*k, l, i*) contributes to the *f_k,l_* entry of the SFS. More specifically, *f_k,l_* is proportional to the total probability (from *t* = 0 to infinity) of lineages (*k, l*, 1) and (*k, l*, 2).

Assume that in the matrix form the equation has the form

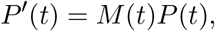

where *P* is the vector of probabilities of states, and M is a transition matrix depending on coalescence and migration rates. In order to calculate SFS, we need to compute the corresponding integrals 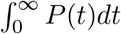 of the time spent in each of the states. We assume that the local effective population sizes and migration rates are picewise constant, hence *M* is piecewise constant too. On each time interval [t_0_, t_1_] the solution of the matrix equation is *P*(*t*) = exp(*Mt*)*P*(*t*_0_), and the integral

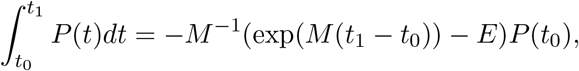

where exp is the matrix exponent and *E* is the identity matrix.

### A.2 Simulations

Simulations were done with the software *ms* [Hudson, 2002] and using GNU parallel [Tange, 2011] to run replicates and explore parameter values. First, we simulated two types of populations: *Population 1* remained with constant intermediate size, while *population 2* underwent a bottleneck followed by expansion, similar to the scenario simulated in Figure 2 of Li and Durbin [2011]. The exact ms command line for simulating size changes of populations 1 and 2 is the following: “4 100 -t 15000 -r 1920 30000000 -l -I 2 2 2 -n 1 1.5 -n 2 3.0 -en 0.025 2 0.2 -en 0.175 2 1.5 -ej 0.625 2 1 -eN 3 3”.

We added continuous migration using the *-em* flag in ms, and pulse migration was included using *-es* to split the receiver population into a third population, followed by *-ej* to merge the third population into the donor population at the exact same time.

To simulate a single panmictic population with the same historical ordinary effective population size as population 2 (used for Figure 4C,E), we used the following ms command line: “2 10000 -T -t 15000 -r 1920 30000000 -l -eN 0.0 3.233 -eN 0.01 3.69 -eN 0.02 3.996 -eN 0.025 0.311 -eN 0.03 0.359 -eN 0.04 0.436 -eN 0.05 0.531 -eN 0.06 0.644 -eN 0.07 0.777 -eN 0.08 0.926 -eN 0.09 1.088 -eN 0.1 1.369 -eN 0.125 1.723 -eN 0.15 1.928 -eN 0.175 2.274 -eN 0.2 2.049 -eN 0.25 1.859 -eN 0.3 1.703 -eN 0.4 1.597 -eN 0.5 1.545 -eN 0.625 1.5 -eN 3.0 3.0”.

We also simulated past population size changes that closely approximate those inferred by PSMC for human populations. An approximation of the **Han-French** effective population size trajectories, was simulated using the command: “4 1000 -t 1500 -r 192 3000000 -l -I 2 2 2 -n 1 6.3 -n 2 2.4 -ej x 2 1 -eN 0.0225 0.3 -eN 0.15 2.7 -eN 0.5 1.3 -eN 2.5 2.9 -eN 5 2.7”, using the following values of the split times (x): 0.0225 (900 generations, the end of the bottleneck), 0.043 (1714 generations, the split time inferred by MiSTI, within the bottleneck), 0.15 (6000 generations, right before the bottleneck) and 0.575 (23000 generations, before the expansion that precedes the main bottleneck) .

An approximation of the **San-Dinka** effective population size trajectories, was simulated using the command: “4 1000 -t 1500 -r 192 3000000 -l -I 2 2 2 -n 1 1 -n 2 1 -ej x 2 1 -en 0.05 1 2 -en 0.11 1 3 -en 0.15 2 2.7 -eN 0.5 1.3 -eN 2.5 2.9 -eN 5 2.7”, using the following values of the split times (x): 0.05 (2000 generations), 0.09 (3728 generations, the split time inferred by MiSTI), 0.21 (8554 generations, the split time inferred by TT), 0.25 (10000 generations) and 0.5 (20000 generations).

An approximation of the **Dinka-Sardinian** effective population size trajectories, was simulated using the command: “4 1000 -t 1500 -r 192 3000000 -l -I 2 2 2 -n 1 1 -n 2 1 -ej x 2 1 -en 0.0225 2 0.3 -eN 0.15 2.7 -eN 0.5 1.3 -eN 2.5 2.9 -eN 5 2.7”, using the following values of the split times (x): 0.0225 (900 generations, the end of the bottleneck), 0.064 (2553 generations, the split time inferred by TT), 0.099 (2963 generations, the split time inferred by MiSTI), 0.25 (10000 generations) and 0.5 (20000 generations).

Times in the ms command lines are given in ms units, *i.e*. generations/(4 × *N*_0_), where *N*_0_ = 10000.

### A.3 Data processing

We applied MiSTI to datasets from human and puma populations. MiSTI takes as input PSMC results for one individual from each population. If estimation of migration rates is desired, a joint site frequency spectrum of both genomes is also required. The joint site frequency spectrum can be generated with ANGSD [Korneliussen et al., 2014], and a Python program is provided with MiSTI to convert ANGSD 2D site frequency spectrum format to MiSTI input format. For both humans and pumas, we applied filters to keep only sites with mapping quality above 30 and coverage between one third and twice the average genomewide coverage. In all analyses of human data, we applied the 1000 Genomes strict accessibility genome mask, and a filter for positions where the ancestral state was conserved among three species of great apes (Chimpanzee, Gorilla and Orangutan). The accessibility mask file can be downloaded from http://ftp.1000genomes.ebi.ac.uk/vol1/ftp/release/20130502/supporting/accessible_genome_masks/20140520.strict_mask.autosomes.bed, and the ancestral state data was downloaded from https://zenodo.org/record/4441887. We ran PSMC with parameters -N25 -t15 -r5 -p “4+25*2+4+6” for both species. For humans, we used a mutation rate of 1.25 × 10^-8^ per base pair per generation, and generation time of 29 years. For pumas, we used mutation rate of 5 × 10^-9^ per base pair per generation, and generation time of 5 years [Saremi et al., 2019].

For the analysis of human data, we downloaded a modern European genome (in bam format) from the CEPH/UTAH (CEU) population from the European Nucleotide Archive (ENA), accession number ERR194158. Neanderthal tracts specific for that individual were obtained from Steinrücken et al. [2018]. The Neanderthal bam file was downloaded from http://cdna.eva.mpg.de/neandertal/altai/AltaiNeandertal/bam/. Bam files from one Han Chinese sample (HGDP00778), one French sample (HGDP00521), one San sample (HGDP01029) and one Dinka sample (DNK02) and one Sardinian sample (HGDP00665) were downloaded from http://cdna.eva.mpg.de/denisova/BAM/human/. When inferring migration from real data, we allowed it to start from the 4th PSMC interval and going until the split time, to avoid using the first PSMC intervals that have a lot of uncertainty.

We ran PSMC on the CEU genome before and after masking its tracts of Neanderthal ancestry, and we used MiSTI to correct the PSMC of the unmasked CEU genome, assuming 1.5% and 3% of pulse admixture from Neanderthals. The split time was set to 662 kya, considering that the average archaic-modern human split time inferred in Prüfer et al. [2014] is 570 thousand years, with a mutation rate of 5 × 10^-10^ per base pair per year, and adjusting for the mutation rate we use here (which translates to 4.3 × 10^-10^ per year). The pulse migration time was set to 60 kya, and the sample age was set to 50 kya, using MiSTI’s *–sdate* parameter.

For the analysis of puma data, we obtained bam files and masks for runs of homozygosity (due to recent inbreeding) from the authors of Saremi et al. [2019]. We focused on one sample from Florida (EVG21) that showed an inflated PSMC trajectory in Saremi et al. [2019], likely due to its known history of admixture with Central American pumas, and another sample from Florida that does not have Central American ancestry (CYP47). We masked the runs of homozygosity from these genomes and we ran PSMC on them. We used MiSTI to correct the inferred effective population size trajectories for a plausible scenario of continuous migration and recent pulse admixture from Central America to Florida, based on the known history of this species. Saremi et al. [2019] inferred that the split time between the Florida pumas and Brazilian pumas was 300 thousand years ago. We have assumed a more recent split time of 200 thousand years between the Florida pumas and the Central American pumas that were ancestors of EVG21, which is the time when PSMC trajectories of CYP47 and EVG21 diverge. The resulting trajectory of the admixed individual, after correcting for its Florida ancestry component, is a putative effective population size trajectory for Central American pumas, that were not sampled.

### A.4 Running MiSTI

To run MiSTI with parameter optimization, we recommend starting with inference of split times without migration. Once the best split time for this model (T*) is found, the user can optimize the migration rates in each direction under split times equal to or larger than T*. The user can run the migration rate optimization from different starting values, and we recommend using gnu-parallel [Tange, 2011] to provide the starting values to MiSTI.

### A.5 Time discretization

MiSTI merges time points from two PSMC files, so that effective population size of both populations are constant on each time interval. This discretization is the principal time scale for MiSTI, and the default search of the split time is performed at the nodes of this discretization. If more precision is needed, one may use -`d N` key to split all the intervals in the search range into *N* equal parts.

### A.6 Effective population size before split

Of course, when we choose a split time point, the estimated values for the effective population size before the split do not necessary coincide. So, we need to find a consensus effective population size from two estimates. The consensus effective population size before the split time is computed so that the expected number of coalescences for two haplotypes be the same as the sum of expected number of coalescence for the first and for the second genomes. More formally, let *P*_1_ and *P*_2_ be the probabilities of lineages sampled from populations 1 and 2 respectively not to coalesce before given time interval based on their corresponding distributions of effective population size. The probability not to coalesce within the time interval of length *t* and with estimated effective population sizes *N*_1_ (from the first genome) and *N*_2_ (from the second genome) is

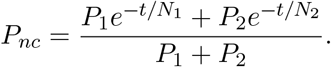

The consensus value for effective population size *N* at this time interval is

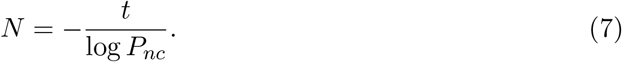

## B Limitations of MiSTI: an analysis of likelihood surfaces

### B.1 The demographic history of a recently admixed puma

In Florida, USA, there is a population of pumas (or mountain lions, *Puma concolor*) known as the Florida Panther. These pumas have a series of morphological signs of inbreeding depression. Interestingly, in the Everglades National Park (EVG), a population does not present these typical signs, likely because of an influx of genetic diversity brought by the introduction of individuals with Central American ancestry O’Brien et al. [1990]. The PSMC trajectory from one of these individuals (EVG21) shows larger effective population sizes than other Florida panthers, as expected due to its admixed ancestry Saremi et al. [2019].

We have corrected the PSMC trajectory of this admixed individual with MiSTI, modeling it is a Central American puma that received admixture from a Florida Panther. We use a sample from the Big Cypress National Preserve (CYP47) as the unadmixed Florida Panther. This population is partially isolated from the EVG population and does not show inflated effective population size like EVG21.

Even though we do not have data from Central American pumas, by using MiSTI to correct the PSMC trajectory of EVG21 for the Florida admixture component (CYP47), we aimed to recover the effective population size trajectory of the unsampled Central American puma.

When we model a split time of 200ky between Central American and Florida panthers, and a single pulse admixture event very close to the present, we can fit a pulse of 0.30 from Florida to Central American. Under this model, we infer that the Central American population had more constant effective population size through time than other puma populations (Figure 9).

**Figure 9:**
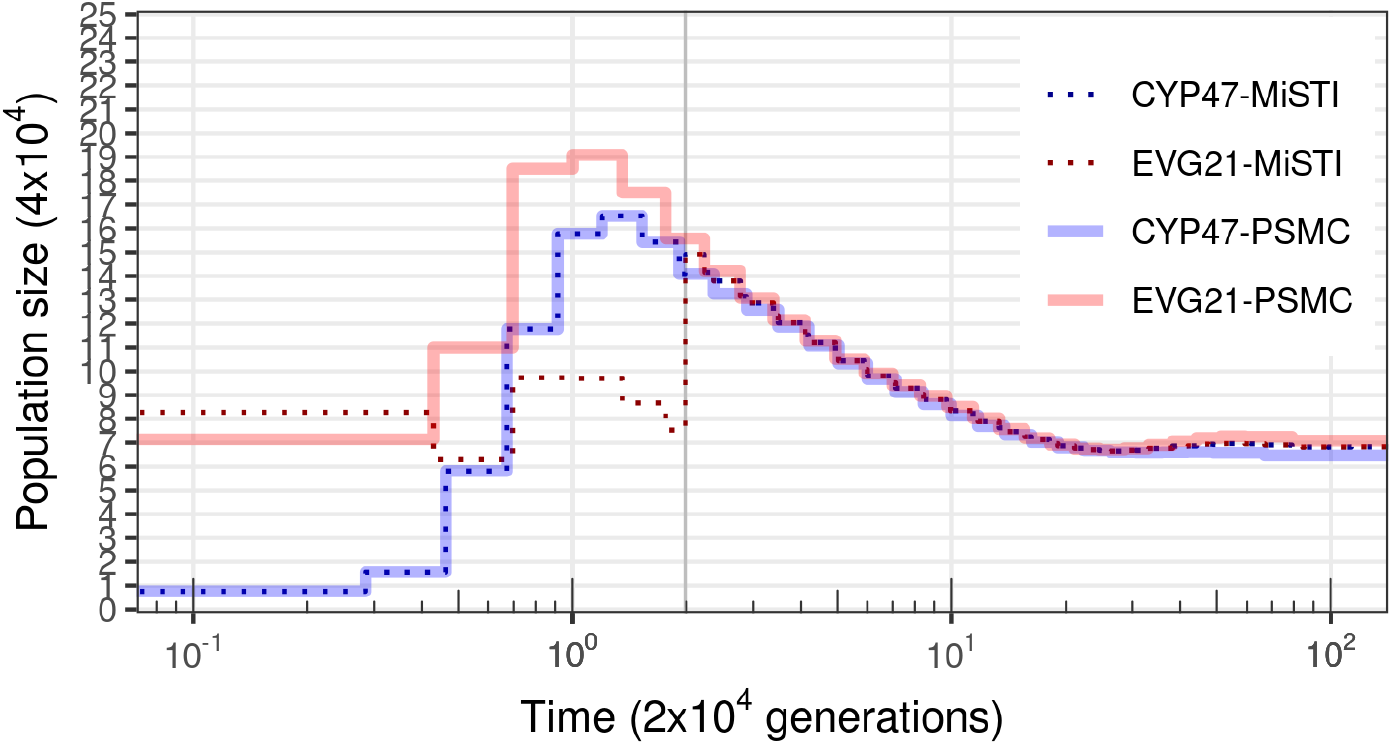
MiSTI correction of the PSMC curve of the admixed puma sample EVG21 with a single pulse of 0.30 admixture from CYP47 at the most recent time interval. Split time (200 thousand years) is indicated by a vertical gray bar.

The range of *Puma concolor* used to be connected from East to West of North America until a great population decline started in the 1800s, when these animals suffered extreme habitat loss and were hunted almost to extinction. Therefore, it is likely that there has been some degree of continuous migration between Florida Panthers and Central American pumas until historical times. Allowing for continuous migration, the model fits the data better (llh=-131360 instead of llh= −172712). This model includes continuous migration (migration rate from Central America to Florida of 1.0, and 0.65 in the other direction) and the same 0.30 recent pulse migration from Florida to Central America. Under this model, we also infer that the Central American puma population had relatively constant effective population size through time, and that both populations had smaller local effective population size than the ancestral effective population size inferred by PSMC (Figure 10). We note that the original PSMC effective population sizes are very high (over 600 thousand individuals), and likely unrealistic for a population of large carnivores. This is likely a consequence of some degree of population structure and continuous migration across the range of pumas until recent times.

**Figure 10:**
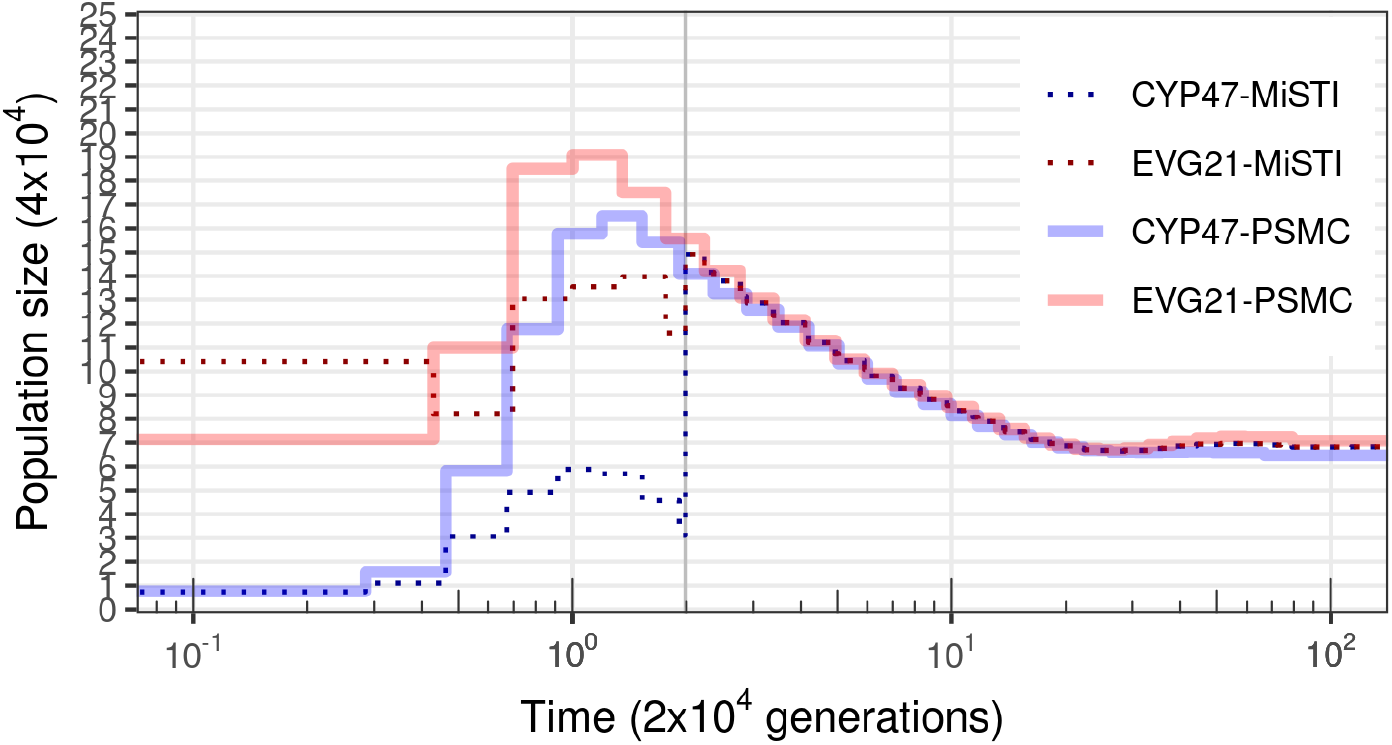
MiSTI correction of the PSMC curve of the admixed puma sample EVG21 with continuous migration allowed since the split time, and a single pulse of 0.30 admixture from CYP47 at the most recent time interval. Split time (200 thousand years) is indicated by a vertical gray bar.

We note, however, that the composite likelihood surface for the split time and pulse of migration used as a model for the Pumas is not smooth (Figure 11). The empty area of the composite likelihood surface indicates rates of continuous migration that are incompatible with the PSMC trajectories. For those values of migration rates, the corrected local effective population size would become negative, which is nonsensical. The fact that the highest composite likelihoods are at the border of the likelihood surface and next to these incompatible values of migration rates indicates that the data does not fit well with the MiSTI model of pulse and continuous migration we used. This uneven composite likelihood surface could also be due to some issue in the PSMC inference step. One possible source of problems is that there is large uncertainty in the PSMC inference at the most recent time intervals, which is when we model the pulse of migration in the Pumas.

**Figure 11:**
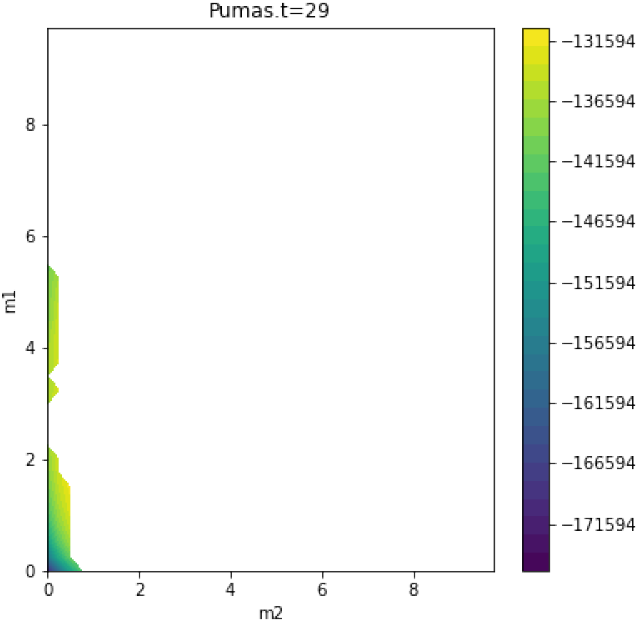
Composite likelihood surface for the model of Florida panther split time 200 kya, pulse of migration at the most recent time interval, and a range of continuous migration rates.

### B.2 Likelihood surfaces of inferred models of split and migration between pairs of human populations

Here we show the composite likelihood surfaces for a range of migration rates under the best inference of split time for pairs of human populations discussed in the main text. These composite likelihood surfaces differ from the puma case above in that their peak is not near the steep drop in composite likelihood.

## C Simulations of the San-Dinka split time with migration

Here we show simulations of the best inference of split time and migration rates for the San-Dinka pair. We did ten replicate simulations with the effective population size trajectories inferred from the data with PSMC, and with split time 3729 generations ago, and migration rate of 2.5 from Dinka to San (Table 2). We applied both the TT method and MiSTI to infer split times in each simulation. The TT method largely overestimated the split time in all cases, while MiSTI underestimated the split time when no migration is allowed in the model (left most column, Figure 15). When migration in the direction simulated (from Dinka to San) is allowed, MiSTI estimates split times closer to the simulated values, and migration rates are also estimated in the correct direction (columns 2 and 4, Figure 15). When migration is only allowed in the direction opposite to the simulated, it is largely overestimated, and the split time is underestimated (third column of Figure 15).

**Figure 12:**
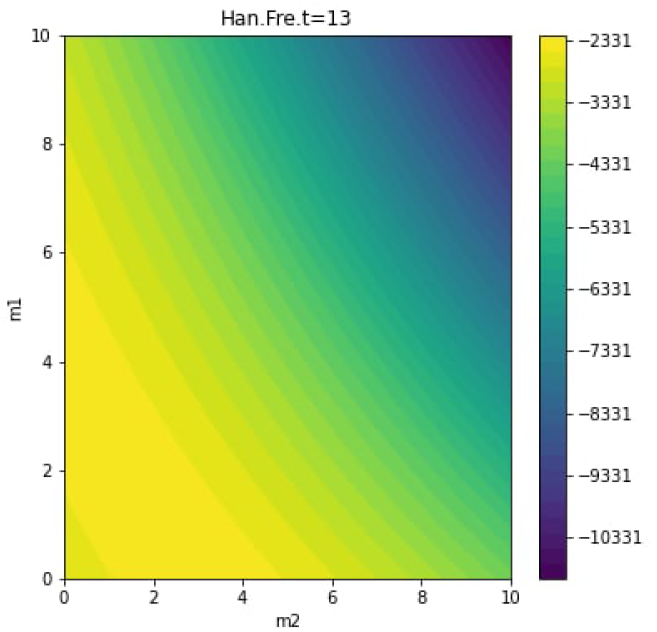
Composite likelihood surface from MiSTI for the inferred split time between Han and French (1505 generations ago), for different values of migration rates.

**Figure 13:**
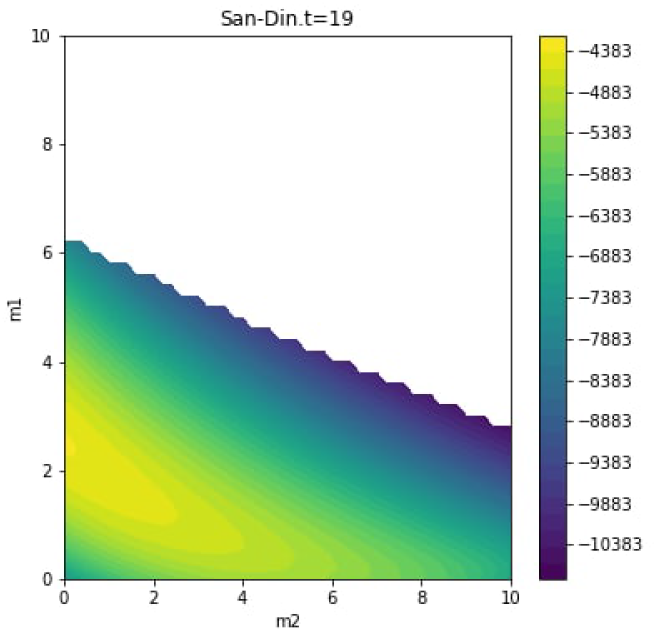
Composite likelihood surface from MiSTI for the inferred split time between San and Dinka (3729 generations ago), for different values of migration rates.

**Figure 14:**
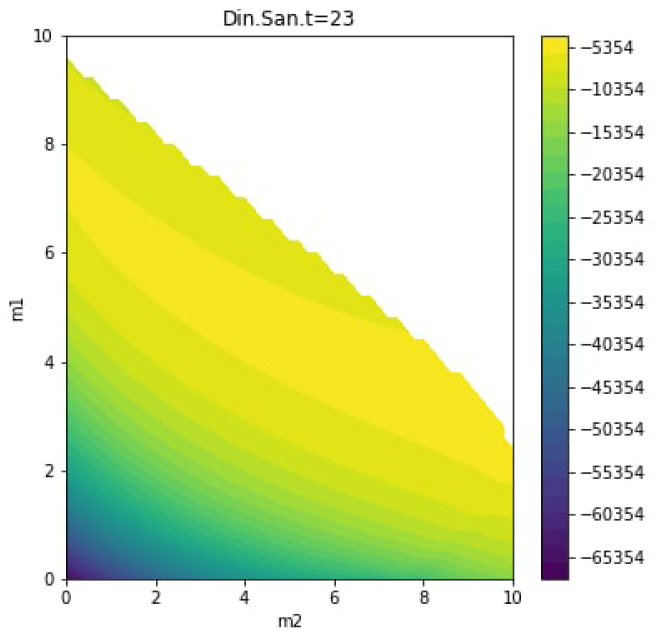
Composite likelihood surface from MiSTI for the inferred split time between Dinka and Sardinian (3963 generations ago), for different values of migration rates.

**Figure 15:**
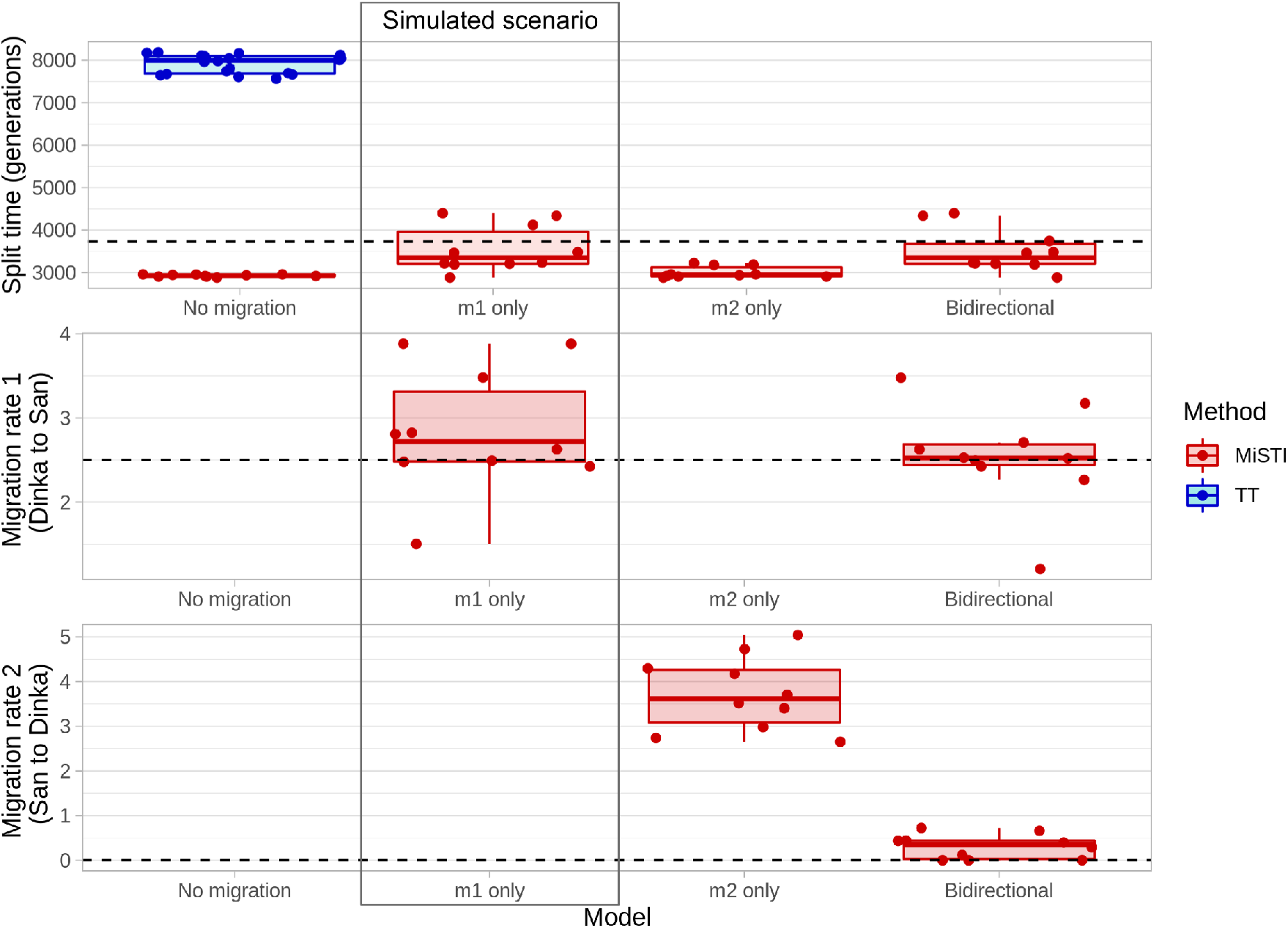
Ten simulations of the split time and migration rates between San and Dinka inferred by MiSTI (split 3729 generations ago, m1=2.5 and m2=0, shown in the main text Table 2). In the top panel, we show split times inferred using the TT method and MiSTI. Middle and bottom panels shows values of inferred migration rates. In MiSTI, we inferred split times and migration rates for 4 models: no migration, m1 only, m2 only and bidirectional migration.

## References

Brenna M. Henn, Luigi Luca Cavalli-Sforza, and Marcus W Feldman. The great human expansion. Proceedings of the National Academy of Sciences of the United States of America, 109(44):17758–64, 10 2012. ISSN 1091-6490. doi: 10.1073/pnas.1212380109. URL http://www.pubmedcentral.nih.gov/articlerender.fcgi?artid=3497766&tool=pmcentrez&rendertype=abstract.

Sewall Wright. Evolution in Mendelian Populations. Genetics, 16(2):97–159, 3 1931. ISSN 0016-6731. URL http://www.pubmedcentral.nih.gov/articlerender.fcgi?artid=1201091%7B&%7Dtool=pmcentrez%7B%7Drendertype=abstract http://www.pubmedcentral.nih.gov/articlerender.fcgi?artid=1201091&tool=pmcentrez&rendertype=abstract.

Daniel L. Hartl and Andrew G. Clark. Principles of Population Genetics. Sinauer Associates, Sunderland, 4th edition, 2007.

R A Fisher. The Genetical Theory of Natural Selection. Clarendon Press, 1930.

P A P Moran. Random processes in genetics. Mathematical Proceedings of the Cambridge Philosophical Society, 54(1):60–71, 1958. doi: 10.1017/S0305004100033193.

J F C Kingman. On the Genealogy of Large Populations. Journal of Applied Probability, 19:27–43, 1982. URL https://www.jstor.org/stable/3213548.

P Sjödin, I Kaj, S Krone, M Lascoux, and M Nordborg. On the meaning and existence of an effective population size. Genetics, 169(2):1061–70, 2 2005. ISSN 0016-6731. doi: 10.1534/genetics.104.026799. URL http://www.ncbi.nlm.nih.gov/pubmed/15489538 http://www.pubmedcentral.nih.gov/articlerender.fcgi?artid=PMC1449138.

John Wakeley and Ori Sargsyan. Extensions of the coalescent effective population size. Genetics, 181(1):341–5, 1 2009. ISSN 0016-6731. doi: 10.1534/genetics.108.092460. URL http://www.ncbi.nlm.nih.gov/pubmed/19001293 http://www.pubmedcentral.nih.gov/articlerender.fcgi?artid=PMC2621185.

Heng Li and Richard Durbin. Inference of human population history from individual whole-genome sequences. Nature, 475(7357):493–496, 7 2011. ISSN 0028-0836. doi: 10.1038/nature10231. URL http://www.nature.com/articles/nature10231.

Gilean A T McVean and Niall J Cardin. Approximating the coalescent with recombination. Philosophical transactions of the Royal Society of London. Series B, Biological sciences, 360(1459):1387–93, 2005. ISSN 0962-8436. doi: 10.1098/rstb.2005.1673. URL http://www.pubmedcentral.nih.gov/articlerender.fcgi?artid=1569517&tool=pmcentrez&rendertype=abstract.

Asger Hobolth, Ole F Christensen, Thomas Mailund, and Mikkel H Schierup. Genomic Relationships and Speciation Times of Human, Chimpanzee, and Gorilla Inferred from a Coalescent Hidden Markov Model. PLOS Genetics, 3(2):1–11, 2007. doi: 10.1371/journal.pgen.0030007. URL https://doi.org/10.1371/journal.pgen.0030007.

Julien Y. Dutheil, Ganesh Ganapathy, Asger Hobolth, Thomas Mailund, Marcy K. Uyenoyama, and Mikkel H. Schierup. Ancestral population genomics: The coalescent hidden Markov model approach. Genetics, 183(1):259–274, 2009. ISSN 00166731. doi: 10.1534/genetics.109.103010.

Sara Sheehan, Kelley Harris, and Yun S Song. Estimating variable effective population sizes from multiple genomes: a sequentially markov conditional sampling distribution approach. Genetics, 194(3):647–62, 7 2013. ISSN 1943-2631. doi: 10.1534/genetics.112.149096. URL http://www.ncbi.nlm.nih.gov/pubmed/6628982 http://www.pubmedcentral.nih.gov/articlerender.fcgi?artid=PMC1202167 http://www.ncbi.nlm.nih.gov/pubmed/23608192 http://www.pubmedcentral.nih.gov/articlerender.fcgi?artid=PMC3697970.

Stephan Schiffels and Richard Durbin. Inferring human population size and separation history from multiple genome sequences. Nature genetics, 46(8):919–25, 2014. ISSN 1546-1718. doi: 10.1038/ng.3015. URL http://dx.doi.org/10.1038/ng.3015.

Jonathan Terhorst, John A Kamm, and Yun S Song. Robust and scalable inference of population history from hundreds of unphased whole genomes. Nature Genetics, 49(2): 303–309, 2017. ISSN 1061-4036. doi: 10.1038/ng.3748. URL http://www.nature.com/doifinder/10.1038/ng.3748.

Jeffrey P. Spence, Matthias Steinrücken, Jonathan Terhorst, and Yun S. Song. Inference of population history using coalescent HMMs: review and outlook. Current Opinion in Genetics and Development, 53:70–76, 2018. ISSN 18790380. doi: 10.1016/j.gde.2018.07.002.

Rasmus Heller, Lounes Chikhi, and Hans Redlef Siegismund. The Confounding Effect of Population Structure on Bayesian Skyline Plot Inferences of Demographic History. PLoS ONE, 8(5):e62992, 5 2013. ISSN 1932-6203. doi: 10.1371/journal.pone.0062992. URL https://dx.plos.org/10.1371/journal.pone.0062992.

O Mazet, W Rodríguez, S Grusea, S Boitard, and L Chikhi. On the importance of being structured: instantaneous coalescence rates and human evolution—lessons for ancestral population size inference? Heredity, 116(4):362–371, 4 2016. ISSN 0018-067X. doi: 10.1038/hdy.2015.104. URL http://www.nature.com/articles/hdy2015104.

Lounès Chikhi, Willy Rodríguez, Simona Grusea, Patrícia Santos, Simon Boitard, and Olivier Mazet. The IICR (inverse instantaneous coalescence rate) as a summary of genomic diversity: insights into demographic inference and model choice. Heredity, 120:13–24, 1 2018. ISSN 0018-067X. doi: 10.1038/s41437-017-0005-6. URL http://www.nature.com/articles/s41437-017-0005-6.

Jinliang Wang and Michael C. Whitlock. Estimating effective population size and migration rates from genetic samples over space and time. Genetics, 163(1):429–446, 2003. ISSN 00166731.

Ke Wang, Iain Mathieson, Jared O’Connell, and Stephan Schiffels. Tracking human population structure through time from whole genome sequences. PLoS Genetics, 16(3):1–24, 2020. ISSN 15537404. doi: 10.1371/journal.pgen.1008552. URL http://dx.doi.org/10.1371/journal.pgen.1008552.

Shiya Song, Elzbieta Sliwerska, Sarah Emery, and Jeffrey M. Kidd. Modeling human population separation history using physically phased genomes. Genetics, 205(1):385–395, 2017. ISSN 19432631. doi: 10.1534/genetics.116.192963.

Armando Arredondo, Beatriz Mourato, Khoa Nguyen, Simon Boitard, Willy Rodríguez, Olivier Mazet, and Lounes Chikhi. Inferring number of populations and changes in connectivity under the n-island model. Heredity, 126(6):896–912, 2021. ISSN 1365-2540. doi: 10.1038/s41437-021-00426-9. URL https://doi.org/10.1038/s41437-021-00426-9.

Carina M. Schlebusch, Helena Malmström, Torsten Günter, Per Sjödin, Alecandra Coutinho, Hanna Edlund, Arielle R. Munters, Mário Vicente, Maryna Steyn, Himla Soodyall, Marlize Lombard, and Mattias Jakobsson. Southern African ancient genomes estimate modern human divergence to 350,000 to 260,000 years ago. Science, 358:652–655, 2017.

Per Sjödin, James McKenna, and Mattias Jakobsson. Estimating divergence times from DNA sequences. Genetics, 217(4), 2021. ISSN 19432631. doi: 10.1093/genetics/iyab008.

Montgomery Slatkin. Testine neutrality in subdivided populations. Genetics, 100(3):533–545, 1982. ISSN 1943-2631. doi: 10.1093/genetics/100.3.533. URL https://doi.org/10.1093/genetics/100.3.533.

Montgomery Slatkin. Gene Flow and the Geographic Structure of Natural Populations. Science, 236(4803):787–792, 1987. doi: 10.1126/science.3576198. URL https://www.science.org/doi/abs/10.1126/science.3576198.

M Notohara. The coalescent and the genealogical process in geographically structured population. Journal of Mathematical Biology, 29(1):59–75, 1990. ISSN 1432-1416. doi: 10.1007/BF00173909. URL https://doi.org/10.1007/BF00173909.

Hilde M Wilkinson-Herbots. Genealogy and subpopulation differentiation under various models of population structure. Journal of Mathematical Biology, 37(6):535–585, 1998. ISSN 1432-1416. doi: 10.1007/s002850050140. URL https://doi.org/10.1007/s002850050140.

R. R. Hudson. Generating samples under a Wright-Fisher neutral model of genetic variation. Bioinformatics, 18(2):337–338, 2002. ISSN 1367-4803. doi: 10.1093/bioinformatics/18.2.337. URL https://academic.oup.com/bioinformatics/article-lookup/doi/10.1093/bioinformatics/18.2.337.

Kay Prüfer, Fernando Racimo, Nick J Patterson, Flora Jay, Sriram Sankararaman, Susanna Sawyer, Anja Heinze, Gabriel Renaud, Peter H Sudmant, Cesare de Filippo, Heng Li, Swapan Mallick, Michael Dannemann, Qiaomei Fu, Martin Kircher, Martin Kuhlwilm, Michael Lachmann, Matthias Meyer, Matthias Ongyerth, Michael Siebauer, Christoph Theunert, Arti Tandon, Priya Moorjani, Joseph K. Pickrell, James C Mullikin, Samuel H Vohr, Richard E Green, Ines Hellmann, Philip L F Johnson, Hélène Blanche, Howard Cann, Jacob O Kitzman, Jay Shendure, Evan E Eichler, Ed S Lein, Trygve E Bakken, Liubov V Golovanova, Vladimir B Doronichev, Michael V Shunkov, Anatoli P Derevianko, Bence Viola, Montgomery Slatkin, David E Reich, Janet Kelso, and Svante Pääbo. The complete genome sequence of a Neanderthal from the Altai Mountains. Nature, 505(7481):43–9, 2014. ISSN 1476-4687. doi: 10.1038/nature12886. URL http://www.pubmedcentral.nih.gov/articlerender.fcgi?artid=4031459&tool=pmcentrez&rendertype=abstract.

Fernando A. Villanea and Joshua G. Schraiber. Multiple episodes of interbreeding between Neanderthal and modern humans. Nature Ecology & Evolution, 3(1):39–44, 1 2019. ISSN 2397-334X. doi: 10.1038/s41559-018-0735-8. URL http://www.nature.com/articles/s41559-018-0735-8.

Matthias Steinrücken, Jeffrey P. Spence, John A. Kamm, Emilia Wieczorek, and Yun S. Song. Model-based detection and analysis of introgressed Neanderthal ancestry in modern humans. Molecular Ecology, 27(19):3873–3888, 10 2018. ISSN 09621083. doi: 10.1111/mec.14565. URL http://doi.wiley.com/10.1111/mec.14565.

Richard E Green, Johannes Krause, Adrian W Briggs, Tomislav Maricic, Udo Stenzel, Martin Kircher, Nick J Patterson, Heng Li, Weiwei Zhai, Markus Hsi-Yang Fritz, Nancy F Hansen, Eric Y. Durand, Anna-sapfo Malaspinas, Jeffrey D. Jensen, Tomas Marques-Bonet, Can Alkan, Kay Prüfer, Matthias Meyer, Hernán A Burbano, Jeffrey M Good, Rigo Schultz, Ayinuer Aximu-Petri, Anne Butthof, Barbara Höber, Barbara Höffner, Madlen Siegemund, Antje Weihmann, Chad Nusbaum, Eric S Lander, Carsten Russ, Nathaniel Novod, Jason Affourtit, Michael Egholm, Christine Verna, Pavao Rudan, Dejana Brajkovic, Zeljko Kucan, Ivan Gušoronichev, Liubov V Golovanova, Carles Lalueza-Fox, Marco de la Rasilla, Javier Fortea, Antonio Rosas, Ralf W Schmitz, Philip L F Johnson, Evan E Eichler, Daniel Falush, Ewan Birney, James C Mullikin, Montgomery Slatkin, Rasmus Nielsen, Janet Kelso, Michael Lachmann, David E Reich, and Svante Pääbo. A Draft Sequence of the Neandertal Genome. Science, 328(5979): 710–722, 2010.

Wolfgang Haak, Iosif Lazaridis, Nick J Patterson, Nadin Rohland, Swapan Mallick, Bastien Llamas, Guido Brandt, Susanne Nordenfelt, Éadaoin Harney, Kristin Stewardson, Qiaomei Fu, Alissa Mittnik, Eszter Bánffy, Christos Economou, Michael Francken, Susanne Friederich, Rafael Garrido Pena, Fredrik Hallgren, Valery Khartanovich, Aleksandr Khokhlov, Michael Kunst, Pavel Kuznetsov, Harald Meller, Oleg Mochalov, Vayacheslav Moiseyev, Nicole Nicklisch, Sandra L. Pichler, Roberto Risch, Manuel A. Rojo Guerra, Christina Roth, Anna Szécséenyi-Nagy, Joachim Wahl, Matthias Meyer, Johannes Krause, Dorcas Brown, David Anthony, Alan Cooper, Kurt Werner Alt, and David Reich. Massive migration from the steppe was a source for Indo-European languages in Europe. Nature, 522(7555):207–211, 2015. ISSN 0028-0836. doi: 10.1038/nature14317. URL http://www.nature.com/doifinder/10.1038/nature14317.

Priya Moorjani, Nick J Patterson, Joel N. Hirschhorn, Alon Keinan, Li Hao, Gil Atzmon, Edward Burns, Harry Ostrer, Alkes L. Price, and David E Reich. The history of african gene flow into Southern Europeans, Levantines, and Jews. PLoS Genetics, 7(4), 2011. ISSN 15537390. doi: 10.1371/journal.pgen.1001373.

Luca Pagani, Daniel John Lawson, Evelyn Jagoda, Alexander Mörseburg, Anders Eriksson, Mario Mitt, Florian Clemente, Georgi Hudjashov, Michael DeGiorgio, Lauri Saag, Jeffrey D Wall, Alexia Cardona, Reedik Mägi, Melissa A Wilson Sayres, Sarah Kaewert, Charlotte Inchley, Christiana L Scheib, Mari Järve, Monika Karmin, Guy S Jacobs, Tiago Antao, Florin Mircea Iliescu, Alena Kushniarevich, Qasim Ayub, Chris Tyler-Smith, Yali Xue, Bayazit Yunusbayev, Kristiina Tambets, Chandana Basu Mallick, Lehti Saag, Elvira Pocheshkhova, George Andriadze, Craig Muller, Michael C Westaway, David M Lambert, Grigor Zoraqi, Shahlo Turdikulova, Dilbar Dalimova, Zhaxylyk Sabitov, Gazi Nurun Nahar Sultana, Joseph Lachance, Sarah Tishkoff, Kuvat Momynaliev, Jainagul Isakova, Larisa D Damba, Marina Gubina, Pagbajabyn Nymadawa, Irina Evseeva, Lubov Atramentova, Olga Utevska, François-Xavier Ricaut, Nicolas Brucato, Herawati Sudoyo, Thierry Letellier, Murray P Cox, Nikolay A Barashkov, Vedrana Skaro, Lejla Mulahasanović, Dragan Primorac, Hovhannes Sahakyan, Maru Mormina, Christina A Eichstaedt, Daria V Lichman, Syafiq Abdullah, Gyaneshwer Chaubey, Joseph T S Wee, Evelin Mihailov, Alexandra Karunas, Sergei Litvinov, Rita Khusainova, Natalya Ekomasova, Vita Akhmetova, Irina Khidiyatova, Damir Marjanović, Levon Yepiskoposyan, Doron M Behar, Elena Balanovska, Andres Metspalu, Miroslava Derenko, Boris Malyarchuk, Mikhail Voevoda, Sardana A Fedorova, Ludmila P Osipova, Marta Mirazóon Lahr, Pascale Gerbault, Matthew Leavesley, Andrea Bamberg Migliano, Michael Petraglia, Oleg Balanovsky, Elza K Khusnutdinova, Ene Metspalu, Mark G Thomas, Andrea Manica, Rasmus Nielsen, Richard Villems, Eske Willerslev, Toomas Kivisild, and Mait Metspalu. Genomic analyses inform on migration events during the peopling of Eurasia. Nature, 538(7624):238–242, 2016. ISSN 1476-4687. doi: 10.1038/nature19792. URL https://doi.org/10.1038/nature19792.

Shaohua Fan, Derek E Kelly, Marcia H Beltrame, Matthew E B Hansen, Swapan Mallick, Alessia Ranciaro, Jibril Hirbo, Simon Thompson, William Beggs, Thomas Nyambo, Sabah A Omar, Dawit Wolde Meskel, Gurja Belay, Alain Froment, Nick Patterson, David Reich, and Sarah A Tishkoff. African evolutionary history inferred from whole genome sequence data of 44 indigenous African populations. Genome Biology, 20(1):82, 2019. ISSN 1474-760X. doi: 10.1186/s13059-019-1679-2. URL https://doi.org/10.1186/s13059-019-1679-2.

Jack Kamm, Jonathan Terhorst, Richard Durbin, and Yun S Song. Efficiently inferring the demographic history of many populations with allele count data. Journal of the American Statistical Association, 115(531):1472–1487, 2020. ISSN 0162-1459 (Print). doi: 10.1080/01621459.2019.1635482.

Anders Bergström, Chris Stringer, Mateja Hajdinjak, Eleanor M.L. Scerri, and Pontus Skoglund. Origins of modern human ancestry. Nature, 590(7845):229–237, 2021. ISSN 14764687. doi: 10.1038/s41586-021-03244-5. URL http://dx.doi.org/10.1038/s41586-021-03244-5.

O Tange. GNU Parallel - The Command-Line Power Tool. ;login: The USENIX Magazine, 36(1):42–47, 2 2011. doi: 10.5281/zenodo.16303. URL http://www.gnu.org/s/parallel.

Thorfinn S Korneliussen, Anders Albrechtsen, and Rasmus Nielsen. ANGSD: analysis of next generation sequencing data. BMC bioinformatics, 15(1):356, 2014.

Nedda F Saremi, Megan A Supple, Ashley Byrne, James A Cahill, Luiz Lehmann Coutinho, Love Dalén, Henrique V Figueiró, Warren E Johnson, Heather J Milne, Stephen J O’Brien, Brendan O’Connell, David P Onorato, Seth P D Riley, Jeff A Sikich, Daniel R Stahler, Priscilla Marqui Schmidt Villela, Christopher Vollmers, Robert K Wayne, Eduardo Eizirik, Russell B Corbett-Detig, Richard E Green, Christopher C Wilmers, and Beth Shapiro. Puma genomes from North and South America provide insights into the genomic consequences of inbreeding. Nature Communications, 10(1):4769, 2019. ISSN 2041-1723. doi: 10.1038/s41467-019-12741-1. URL https://doi.org/10.1038/s41467-019-12741-1.

Stephen J. O’Brien, Melody E. Roelke, Naoya Yuhki, Karen W. Richards, Warren E. Johnson, William L. Franklin, Allen E. Anderson, Oron L. Bass Jr., Robert C. Belden, and Janice S. Martenson. Genetic introgression within the Florida Panther Felis concolor coryi. National Geographic Research, 6(4):485–494, 1990. doi: 10.1055/s-2008-1040325.

